# Phylogenetic evidence for asparagine to aspartic acid protein editing of N-glycosylated SARS-CoV-2 viral proteins by NGLY1 deglycosylation/deamidation suggests an unusual vaccination strategy

**DOI:** 10.1101/2021.09.11.459891

**Authors:** Gary Ruvkun, Fei Ji, Ruslan I. Sadreyev

**Affiliations:** Department of Molecular Biology Massachusetts General Hospital, Simches Research Building, Boston MA 02114; Department of Genetics, Harvard Medical School, Simches Research Building, Boston MA 02114

**Author notes:** Corresponding author Gary Ruvkun.

## Abstract

Many viral proteins, including multiple SARS-CoV-2 proteins, are secreted via the endoplasmic reticulum, and viral particles are assembled and exported in ER-associated replication compartments. Viral coat proteins such as the SARS-CoV-2 Spike protein are N-glycosylated at NxS/T sites as they enter the ER. N-glycosylated sites in many eukaryotic proteins are deglycosylated by the NGLY1/PNG-1 deglycosylation enzyme which also deamidates the N-glycosylated asparagine to aspartic acid, thus editing the target protein sequence. Proteomic analysis of mammalian cell lines has revealed deamidation of many host N-glycosylated asparagines to aspartic acid by NGLY1/PNG-1 on peptides that are presented by mammalian HLA for immune surveillance. The key client protein for NGLY1/PNG-1 deglycosylation and N to D protein editing was revealed by genetic analysis of *C. elegans* proteasome regulation to be the intact endoplasmic reticulum-transiting SKN-1A transcription factor. Strikingly, an analysis of cancer cell genetic dependencies for growth revealed that the mammalian orthologue of SKN-1A, NRF1 (also called NFE2L1) is required by a highly correlated set of cell lines as NGLY1/PNG-1, supporting that NGLY1/PNG-1 and NRF1 act in the same pathway. NGLY1/PNG-1 edits N-glycosylated asparagines on the intact SKN-1 protein as it is retrieved by ERAD from the ER to in turn activate the transcription of target proteasomal genes. The normal requirement for NGLY1/PNG-1 editing of SKN-1A can be bypassed by a genomic substituion of N to D in four NxS/T N-glycosylation motifs of SKN-1A. Thus NGLY1/PNG-1-mediated N to D protein editing is more than a degradation step for the key client protein for proteasomal homeostasis in *C. elegans* or tumor growth in particular mammalian cell lines, SKN-1A/NRF1. In addition, such N to D substitutions in NxS/T N-glycosylation motifs occur in evolution: N to D substitutions are observed in phylogenetic comparisons of SKN-1A between nematode species that diverged hundreds of millions of years ago or of the vertebrate NRF1 between disparate vertebrates. Genomic N to D mutations bypass the many steps in N-glycosylation in the ER and deglycosylation-based editing of N to D, perhaps based on differences in the competency of divergent species for various N-glycosylation or deglycosylation steps.

We surveyed the N-glycosylation sites in coronavirus proteins for such phylogenetic evidence for N to D protein editing in viral life cycles, and found evidence for preferential N to D residue substitutions in NxS/T N-glycosylation sites in comparisons of the genome sequences of hundreds of coronaviruses. This suggests that viruses use NGLY1/PNG-1 in some hosts, for example humans, to edit particular N-glycosylated residues to aspartic acid, but that in other hosts, often in bats, an N to D substitution mutation in the virus genome is selected. Single nucleotide mutations in Asp or Asn codons can produce viruses with N to D or D to N substitutions that might be selected in different animal hosts from the population of viral variants produced in any previous host. NGLY1/PNG-1 has been implicated in viral immunity in mammalian cell culture, favoring this hypothesis.

Because of the phylogenetic evidence that the NGLY1/PNG-1 editing of protein sequences has functional importance for SKN-1A/NRF1 and viruses, and because most immunization protocols do not address the probable editing and functional importance of N-glycosylated aspargines to aspartic acid in normal viral infections, we suggest that immunization with viral proteins engineered to substitute D at genomically encoded NxS/T sites of N-glycosylated viral proteins that show a high frequency of N to D substitution in viral phylogeny may enhance immunological response to peptide antigens. Such genomically-edited peptides would not require ER-localization for N-glycosylation or other cell compartment localization for NGLY1/PNG-1 N to D protein editing. In addition, such N to D edited protein vaccines could be produced in bacteria since N-glycosylation and deglycosylation which do not occur in bacteria would no longer be required to immunize with a D-substituted peptide. Bacterially-expressed vaccines would be much lower cost and with fewer failure modes than attenuated viral vaccines or recombinant animal viruses produced in chicken eggs, mammalian tissue culture cells, or delivered by mRNA vectors to the patient directly. Because N to D edited peptides are clearly produced by NGLY1/PNG-1, and may be and presented by mammalian HLA, such peptides may more robustly activate T-cell killing or B-cell maturation to mediate more robust viral immunity.

---

Many proteins that are secreted from eukaryotic cells are N-glycosylated at asparagine residues in the endoplasmic reticulum and Golgi apparatus. N-linked glycosylation of asparagine is guided by the NxS/T motif of a client protein and is mediated the oligosaccharyl transferase complex OST at the ER membrane using the lipid glycan carrier N-dolichol. The OST complex interacts with the NxS/T motifs as the N-terminal regions of a nascent peptide emerges into the lumen of the ER from the Sec61 translocon complex from translating ribosomes on the cytoplasmic side of the ER. Disulfides are oxidized during protein folding in the ER after the N-glycosylation; thus, N-glycosylation is an early step in the maturation of secreted proteins after their sorting to secretory ribosomes by their signal sequences. The added glycans of glycosylation increase the solubility of proteins, as well as mediate a wide variety of cytoskeletal and immune binding reactions. N-glycosylation can also hide protein epitopes from immune presentation by HLA or immune selection by B cells in the selection of the particular immunoglobulin rearrangements and epitope variants that best recognize pathogens. N-linked glycosylation is a common feature of viral envelope proteins, used for immune evasion, as well as protein folding, and interaction with host glycan binding partners. The glycosylated envelope proteins can also hide large regions of the protein from humoral or cellular peptide recognition systems. Shedding of viral glycoproteins can also evade the host humoral immune response, as has been observed for the Coronavirus spike protein.

The SARS-CoV-2 coronaviruses encode 29 proteins, many of which pass through the ER and enter the replication organelle, a modified endoplasmic reticulum (Zhang et al, 2020). The Spike protein has 22 predicted N-glycosylation sites many of which have been confirmed by mass spectroscopy or N to A mutagenesis (Zhang et al, 2020). These N-glycosylated asparagines are located in exposed regions of the infective virus particle that mediate the binding to the Ace2 angiotensin converting enzyme receptor and viral entry into host cells (Zhang et al, 2020). In addition to the envelope proteins, additional SARS-CoV-2 proteins are secreted to the ER where many are N-glycosylated and form disulfide bonds in their path towards viral packaging and secretion. A network of membranes connect the ER membrane to the replication organelles where RNA-dependent-RNA polymerase and other virally-encoded RNA polymerase accessory factors copy the positive strand for viral replication and packaging into coat protein and membranes. The viral proteins recruit host ER proteins to generate highly convoluted replication organelles modified from the ER (Knoops et al, 2008). The two large polyprotein precursors are proteolytically processed into RNA replicase subunits that constitute half of the SARS-CoV-2 coding capacity and that localize to replication organelles (Knoops et al., 2008).

## Deglycosylation of N-glycosylated proteins is more than a degradation pathway

Because of the ubiquity and importance of protein N-glycosylation in viral coat proteins for immune evasion and host cell receptor binding, it has been a major focus for decades to dissect mechanisms of viral receptor binding and immune evasion. But deglycosylation of N-glycosylated viral or host proteins has not been a virology focus. Deglycosylation research has focused on a role in protein aggregation, degradation, and disorders of N-glycosylation. The NGLY1/PNG-1 enzyme that deglycosylates and deamidates N-glycosylated asparagines in client proteins of most eukaryotes was discovered two decades ago (Suzuki et al, 2002). Human NGLY1 genetic analysis of disease symptoms of the small number of patients have highlighted a broad developmental and neural function. Mutations in human NGLY1 cause developmental delay, hypotonia, seizure, peripheral neuropathy, and alacrima, a deficit in production of tears (Need et al, 2012). But the molecular pathway that requires NGLY1/PNG-1 deglycosylation was unknown until NGLY1/PNG-1 mutations emerged from *C. elegans* pathway genetic analysis of proteasome capacity, where NGLY1/PNG-1 was shown to be essential for the regulation of proteasome gene expression in response to protein degradation demand in the cell (Lehrbach and Ruvkun, 2016 and 2019). Comprehensive genetic analysis of defective proteasomal degradation pathways in *C. elegans* identified mutations in the NGLY1/PNG-1 deglycosylation enzyme that fail to upregulate proteasome production in the face of inadequate proteasome activity. Loss of function mutations in the NGLY1/PNG-1 deglycosylation enzyme disrupt the deglycosylation of the ER-localized SKN-1A transcription factor to in turn fail to upregulate proteasome gene expression (Figure 1A). Lack of NGLY1/PNG-1 function decouples genes for proteasome structural proteins from transcriptional responses to the detection of insufficient proteasome capacity. This saturation genetics proved that the SKN-1A transcription factor is the key client protein that is deglycosylated by NGLY1/PNG-1. SKN-1A carries a transmembrane domain and is normally ER-localized and then transported to the cytoplasm and degraded by the proteasome via endoplasmic reticulum associated degradation (ERAD). But if proteasome bandwidth for protein degradation is compromised by a proteasome mutation or by a proteasome toxin or by an excess of aggregrated proteins, SKN-1A is not as efficiently degraded and becomes nuclearly-localized to activate the transcription of the multiple classes proteasome genes necessary to generate more proteasome capacity (Lehrbach and Ruvkun, 2016; Lehrbach, Breen, Ruvkun, 2019). Thus the short half-life SKN-1A protein is a key sensor of proteasomal activity so that if proteasome activity is insufficient to degrade SKN-1A, it activates the expression of multiple proteasome subunit genes to generate more proteasomes.

**Figure 1.**
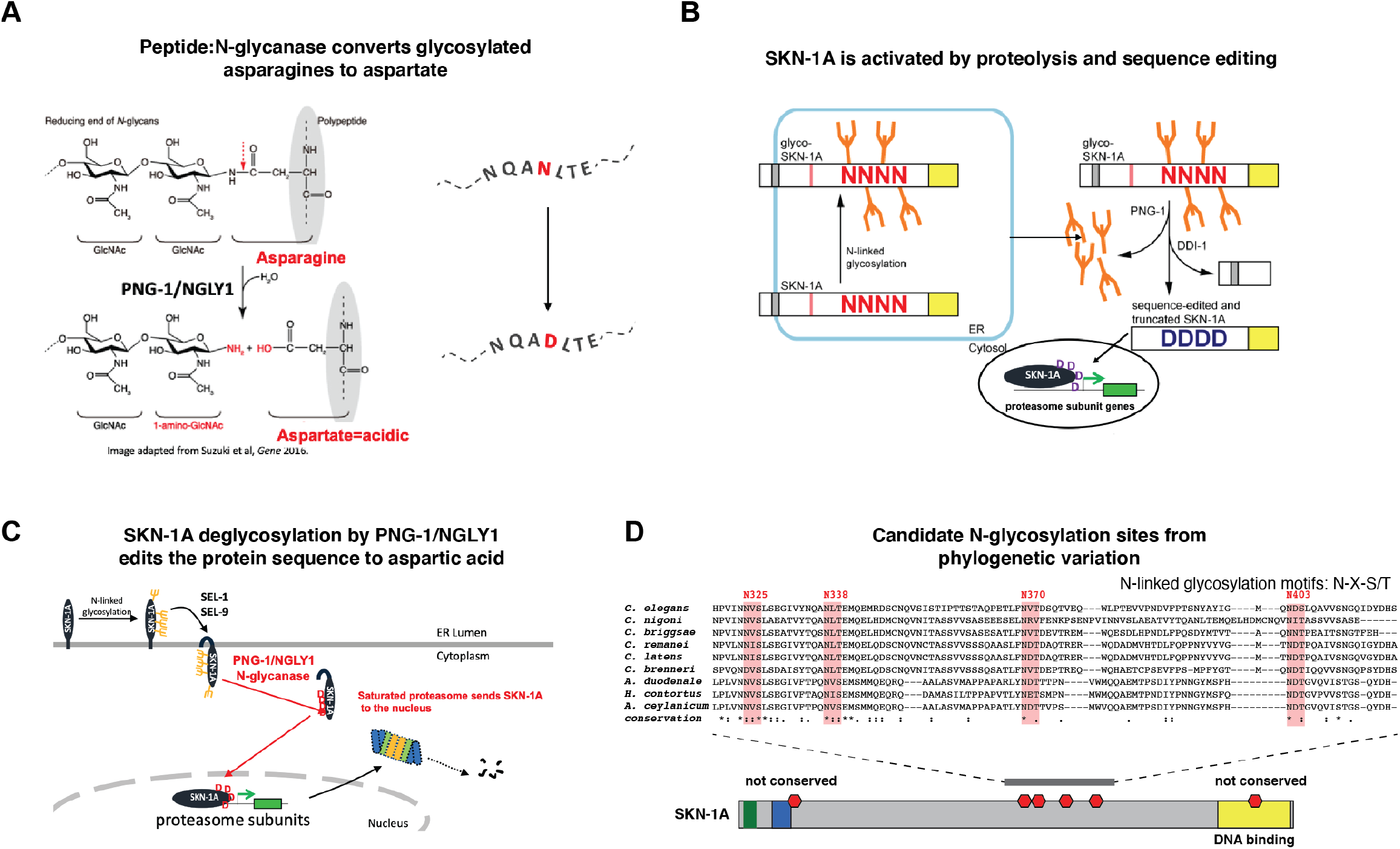
SKN1A protein is a target of asparagine to aspartate sequence editing by PNG/NGLY1 deglycosylase at multiple conserved glycosylation sites. A. Peptide N-glycanase (PNG) catalyzes the conversion of glycosylated asparagine to aspartate (schematic). B. Activation of transcription factor SKN1A through DDI-1 protein cleavage and NGLY1/PNG-1 protein sequence editing. SKN1A is initially glycosylated in endoplasmic reticulum (ER, left). Activation of SKN1A (right) involves PNG-1-mediated deglycosylation and DDI-mediated protein cleavage. C. PNG/NGLY1-mediated deglycosylation of amino acid residues in SKN1A protein results in sequence editing of N-glycosylated asparagines (N) to aspartates (D). D. N-linked glycosylation motifs are conserved in SKN1A proteins across a wide evolutionary range in nematodes. Glycosylation motifs are marked red in the schematic of SKN1A (below). The cluster of four conserved motifs is shown in the multiple sequence alignment across various nematode species (above). Notice that N325 is substituted to D in *C. brenneri*.

*C. elegans png-1* null mutants are viable and healthy unless their proteasome is challenged by a mutation in a proteasomal subunit, by a proteasome inhibitor, or by an abundant aggregated protein, showing that PNG-1 does not have many essential client proteins. Instead, proteasome degradation capacity is the major pathway that requires NGLY1/PNG-1 function in *C. elegans* and most likely in mammals as well.

But how is a deglycosylation enzyme NGLY1/PNG-1 involved in the activation of an ER-localized transcription factor like SKN-1? The mammalian NGLY1 peptide:N-glycanase orthologous to *C. elegans* PNG-1 had been known for decades to remove an amine from the glycosylated asparagine as it deglycosylates and deamidates the client protein to leave behind an aspartic acid residue in the deglycosylated/deamidated client protein asparagine (Suzuki, et al, 2002). But the N-glycosylation field was focused on the removal of carbohydrate modifications of asparagines as a step in protein degradation by the proteasome rather than the N to D protein editing as the key function of NGLY1 deglycosylation. When viewed through the lens of proteasomal protein degradation, this deamination of an asparagine to aspartic acid on the peptides that emerge from the proteasome would be no more interesting than the thousands of other chemical transformations in biochemical degradation pathways. But in the case of the SKN-1A transcription factor activation, the NGLY1/PNG-1 deglycosylation of N-linked glycosylated sites came into an unorthodox focus: the deamidation associated with deglycosylation of N-glycosylated asparagines mediates the editing of the intact SKN-1A protein sequence at those deglycosylated and deamidated asparagines to aspartic acid (an N to D edit--note that this editing by NGLY1/PNG-1 is unrelated to CRISPR DNA editing; it is a very chic protein edit). The *C. elegans* pathway genetics showed that the loss of NGLY1/PNG-1 deglycosylation activity caused the same lack of proteasomal gene activation as loss of the SKN-1A transcription factor.

Thus NGLY1/PNG-1-mediated protein deglycosylation and deamidation **activates** the SKN-1 transcription factor for up-regulation of proteasomal target genes. For example, the glycosylated asparagine could interfere with transcription factor DNA binding or interactions with general transcriptional machinery so that deglycosylation would simply remove the interfering glycan. Or the N to D edit in the SKN-1A protein sequence could be required for this transcription factor interaction, reminiscent of the vogue in “acid blob” transcriptional activation domains 30 years ago (Mitchell 1999). Or the NGLY1/PNG-1 activation of SKN-1A need not have been direct, as is true in any genetic analysis of a multistep pathway. But inspection of the SKN-1<u>A</u> ER-localized isoform, bearing an N-terminal transmembrane domain that specifies secretion to the ER, and N-glycosylation in that compartment, revealed a cluster of four potential N-glycosylation sites that are partially conserved in other nematode species. These potential N-glycosylation sites are not in the bZIP DNA-binding domain located in the C-terminal region. Tantalizingly, in two nematode species (*C. brenneri and C tropicalis*; Figure 1 and Supplementary Table 1), NxS/T at position 325 is substituted to DxS/T, suggesting that N to D protein editing by NGLY1/PNG-1 or a genomic substitution mutation to D is required to activate SKN-1A, not just the removal of a glycan.

To genetically test whether ER localization and N-glycosylation of SKN-1A followed by cytoplasmic deglycosylation and N to D editing is how the SKN-1A transcription factor is actually activated, the *C. elegans* SKN-1A genome sequence was changed to aspartic acid codons at these 4 asparagine codons, a set of four N to D substitution mutations in the genomic *skn-1A* locus. These N to D genome edits activated SKN-1 for the induction of proteasome gene expression even in the absence of NGLY1/PNG-1 deglycosylation, proving that the key client protein for activation of proteasome biogenesis is only SKN-1A and that the 4 asparagines are the key N-glycosylation sites. These N to D genomic substitutions bypassed the normal requirement for NGLY1/PNG-1-mediated protein sequence editing to aspartic acid at the N-glycosylated asparagines of SKN-1. The four N to D mutations bypassed the normal requirement for NGLY1/PNG-1 for resistance to proteasome inhibition because the N to D changes were now in the genome, not via protein editing of N-glycosylated asparagines by NGLY1/PNG-1. This also proved that the N to D editing of SKN-1A by the NGLY1/PNG-1 deglycosylation enzyme is not simply a degradation step on the way to SKN-1A proteolysis. The NGLY1/PNG-1-mediated N to D editing activates SKN-1A as an intact transcription factor that is nuclearly-localized to activate transcription of proteasomal genes (Figure 1B). Proteasomes constitute a major fraction of cellular proteins, so their up-regulation in the face of proteasomal challenges is a major cellular commitment of resources (Figure 1C). The short half-life SKN-1A protein directly detects the decrement in proteasome capacity by its own increased stability in the face of proteasome toxins, mutations, or saturation by protein aggregrates; thus, SKN-1A is both the signal and the transducer of the “need more proteasomes” signal to increase proteasomal gene expression. Single transduction distilled to a detector and transducer in one protein.

One prediction of this N to D editing of ER proteins is that over evolutionary time scales, such edited asparagines might change in the genome sequence to aspartic acid substitutions. Such N to D substitutions would bypass the N to D protein editing dependence on N-glycosylation in the ER and deglycosylation in the cytoplasm, removing the protein from physiological regulation of this protein editing. In fact, comparison of the SKN-1A sequence between nematodes showed that N325 is mutated by a single base change to a D codon in the genome sequence in *C. brenneri*, a species noted for dessication resistance (Sudhaus, 2007). Because N-glycosylation depends on N-dolichol production in the ER, a major lipid, and because deglycosylation of N-glycosylated asparagines uses an oxygen from water for the deamidation of asparagine, there could be selection in particular ecological niches (for example dessicated niches), for such genome sequence substitutions. In addition, given that protein N-glycosylation of viral envelope proteins is a major axis of viral camouflage, there could be selection for activation of such response pathways by genomic editing of host defense protein pathways. SKN-1 mediates many aspects of *C. elegans* defense against pathogens and thus would be subject to such evolutionary arms races. A phylogenetic search for N to D changes in evolution would need to use various time scales determined by the relative selective pressure on the protein domains that are N-glycosylated and by whether the N-glycosylation and deglycosylation to D is part of an evolutionary arms race between for example virus and host. For SKN-1A, the phylogenetic comparison that detected N to D changes was within all nematode species, which are separated by hundreds of millions of years between the parasitic nematodes and *C. elegans*.

## N to D protein editing of the human homologue of *C. elegans* SKN-1A

The human homologue of SKN-1A is NRF1, but also called NFE2L1 or NF2L1 (but is distinct from NRF1 nuclear respiratory factor). Like in most transcription factors, the DNA binding domain, in this case, the 60 amino acid bZip domain, shows a stronger evolutionary conservation and allows the assignment of orthology. The coupling of human NRF1 to NGLY1 deglycosylation was established by molecular studies in the Bertozzi lab (Tomlin et al, 2017). And strikingly, in the cancer genome dependency map of the distinct gene dependencies of hundreds of tumor cell types, there is highly significant similarity of tumor types that are dependent on NRF1, NGLY1, and the aspartic protease DDI2, another proteasomal response pathway gene that emerged from the *C. elegans* genetics (Lehrbach Ruvkun 2016; Wang et al, 2017). For example, 92 out of 990 tumor cell lines are especially sensitive to loss of NGLY1, and 47 cell lines are similarly sensitive to loss of NRF1 transcription factor, and the slow growing cell lines are correlated 0.50 between NRF1 and NGLY1, and 0.65 between DDI2 and NGLY1. These are extraordinarily correlated codependencies. See https://depmap.org/portal/gene/NGLY1?tab=overview. Similarly, NGLY1, NRF1, and DDI2 have emerged from CRISPR screens of mammalian tissue culture cells for hypersensitivity to proteasome inhibition (Kantautas, 2020; Seetharaman, Kantautas, Boone, Andrews, and Moffat, personal communication).

This suggests that exactly as in *C. elegans*, human NRF1 is a major client of NGLY1 protein deglycosylation and deamidation. Human NRF1 like its *C. elegans* SKN-1A orthologue is predicted to have an N-terminal transmembrane domain, to translate at membrane-bound secretory ribosomes, and to localize to the ER. And as its SKN-1A orthologue in *C. elegans*, human NRF1 has been implicated in regulation of proteasome gene expression (Radhakrishnan, 2010). There are multiple predicted NxS/T NRF1 glycosylation sites that are highly conserved between most vertebrates. But the sequence of these N-glycosylation sites begins to diverge in fish. For example, cartilaginous fish are appropriately distant to observe the NxS/T sites diverge to D in the genome sequence, similar to the substitution observed in the *C. brenneri* SKN-1A homologue in nematode phylogeny. For example, an NVS site in human NRF1 diverges to DVS, and another NST diverges to DTS in Chiloscyllium, a shark. A similar set of changes in observed in Poecilia formosa, a fresh water molly fish species (Figure 2A).

**Figure 2.**
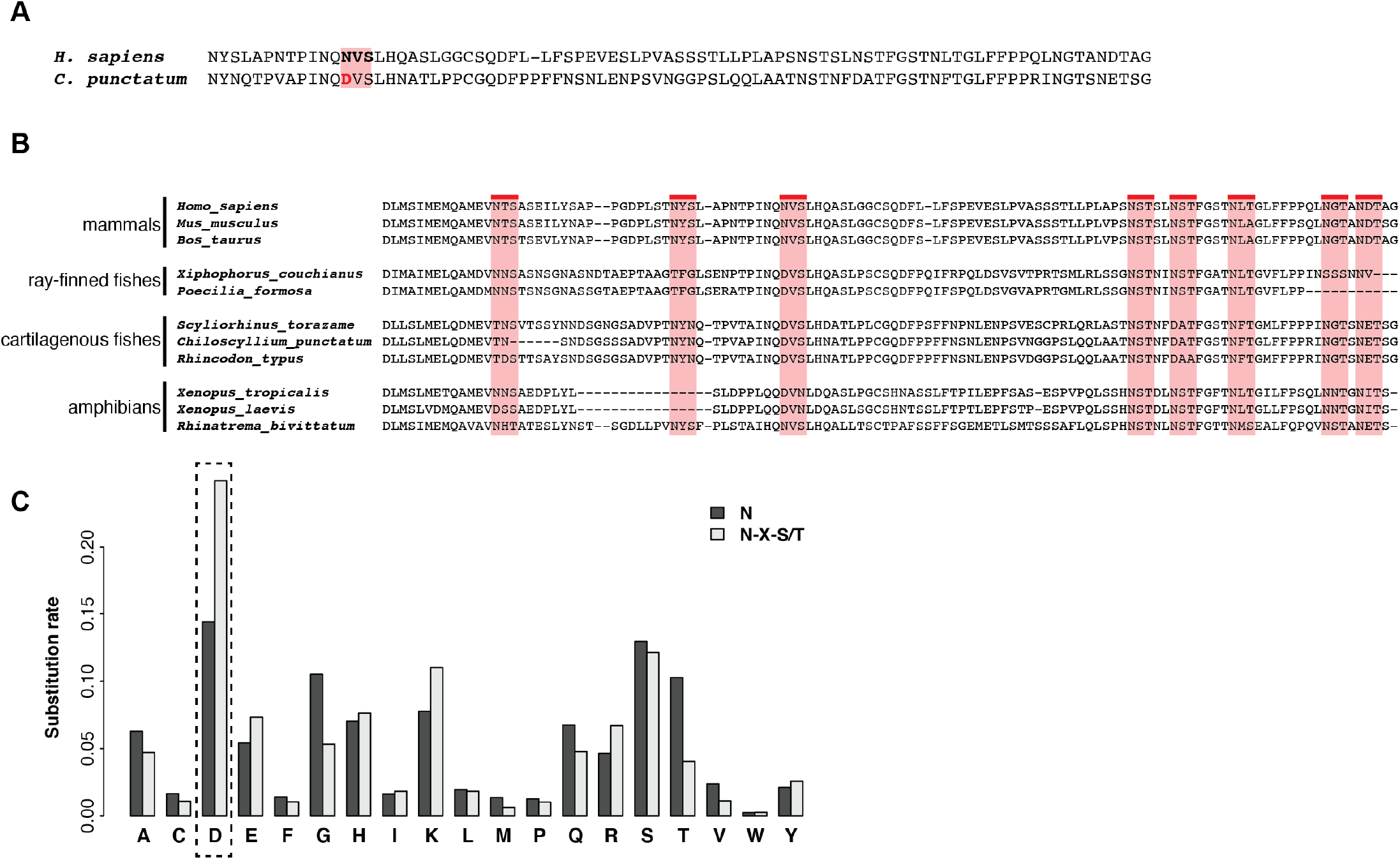
Asparagine in N-linked glycosylation motifs is more frequently mutated to aspartic acid in evolution. A. Example of evolutionary substitutions of asparagine to aspartic acid at N-linked glycosylation motifs among mammalian sequences of NRF1, the human orthologue of SKN1A. B. Multiple sequence alignment of representative NRF1 homologs in mammals, amphibians, and fishes, with human N-linked glycosylation motifs highlighted. C. Among the nematode homologs of *C. elegans* ER proteins, the substitution rate of N to D residue is significantly increased at NxS/T glycosylation motifs compared to general N to D substitution rate in these proteins. The bar plot compares substitution rates of N to other residues across the whole ER proteins (black) to the substitution rates at glycosylation motifs (gray). Notice that N to D but not N to S or N to H are enriched for NxS/T sites.

This sequence analysis of the Nrf1/SKN-1A orthologues supports the view that the N-glycosylation in the ER of this transcription factor followed by NGLY1/PNG-1 deglcosylation with associated deamination is not parochial to *C. elegans* proteasomal homeostasis, but likely to be conserved across animals, including in humans (Figure 2B). In support of NRF1 protein editing of N to D at NxS/T sites activating transcription, N to D substitution mutations in NxS/T sites mediate transcriptional activation of NRF1 target gene expression (Zhang, et al, 2014).

## Mass spectroscopy of mammalian deamidated proteins verifies the NRF1 N to D editing predicted by phylogenetics and shows that they are HLA-presented

Some of the NRF1 N to D edits predicted by phylogenetic comparisons were validated in a recent comprehensive proteomic analysis of protein modifications of HLA-bound peptides by the Purcell and Croft groups at Monash University (Mei et al, 2020). This paper explored peptides carrying particular amino acid modifications that are displayed by the mammalian histocompatibility complex. The presented peptides average nine amino acids long. They analyzed a B-lymphoblastoid cell line stably transfected with different HLA types. These HLA-bound peptides were purified and analyzed by mass spectroscopy, and protein fragments that were mass shifted from the genome sequence predictions by the expected mass and charge of a variety of protein modifications could be assessed. About 500,000 HLA-bound peptides were analyzed: 20K peptides bound to HLA A*01:01, 17K peptides bound to HLA-A*02:01, 15K peptides bound to HLA-A*24:02, 2K peptides bound to HLA-C*04:01, and 429K peptides from published immunopeptide proteomics analyses of melanoma, breast cancer, colon carcinoma, primary fibroblast, and B cell leukemia cells (Mei, et al 2020). They found thousands of peptides that were modified by phosphorylation, deamidation, oxidation, methylation, and other more obscure protein modifications.

Asparagine deamidation is predicted to change the mass of a peptide predicted from the genome sequence by +.98 daltons, the mass difference between –OH and –NH_2_ groups. Asparagine or glutamine deamidations were detected in 2.5% to 7% of the peptides, depending on the HLA type. About ¾ of the 450 deamidations detected in the HLA A*01:01, HLA-A*02:01, HLA-A*24:02, and HLA-C*04:01 complexes were derived from asparagines in the context of an NxS/T consensus N-glycosylation site, strongly suggesting that the deamidation was dependent on previous N-glycosylation and thus likely to be mediated by NGLY1. In fact, their analysis of particular N to D edited peptides in a cell line treated with the NGLY1 inhibitor z-FAD-fmd showed that the deamidation is likely to be NGLY1-mediated (Mei, et al 2020). In addition, many of the peptides showing the N to D deamidation were predicted to derive from ER-localized proteins, where N-glycosylation occurs as the secreted protein enters the ER. The edited aspartic acid was most often localized at position 3 of the nonamer peptide displayed by HLA, suggesting a high level of selection of N to D edited peptides by this highly polymorphic immune display system.

Peptides derived from the SKN-1A orthologue NRF1 protein, with the predicted asparagine to aspartic acid mass change of +.98 daltons, were detected on four NRF1 peptides out of the 450 peptides bound to HLA A*01:01, all of which are located in a predicted NxS/T glycosylation site. Two of those biochemically detected human N to D changes were also detected by the phylogenetic variation of N to D between human and fish species (Figure 2ab).

It may be significant that while the immunoproteomics database of about 500,000 immunopeptides in a variety of cell types and HLA types retrieves almost 100 NRF1 peptides, about 10% of which are deamidated at NxS/T sites, a comparably sized (453K peptides) proteomic database that is not based on HLA presentation, but instead on proteolytic cleavage of proteins isolated from cell lines, retrieved no NRF1 peptides. This suggests that the mammalian HLA peptide presentation pathway and the NRF1 N-glycosylation domain may have coevolved to allow HLA to select NRF1 peptides, perhaps as a surrogate signal of ER stress during a viral infection. The most abundant viral proteins of the envelope are heavily N-glycosylated and many viruses modify the ER to generate membrane bound viral replication and assembly centers. In this way, viral infections disrupt ER activities, for example the N-glycosylation of host proteins, as viral protein synthesis hijacks the ER N-glycosylation systems towards the replication and assembly of viruses (Fischer and Ruvkun, 2020). The normally short half-life NRF1 protein may be stabilized by this proteasome saturation during a viral infection, as proteasomal capacity is saturated by the ERAD retrieval of misfolded host and viral proteins.

It is possible that HLA presentation of N to D edited NRF1 peptides from the N-glycosylated region <u>indicates an unstressed, normally functioning ER</u> in that cell. During a viral infection, NRF1 glycosylation may be less complete, competing with the massive N-glycosylation demands of viral coat proteins, releasing the NRF1 protein to the cytoplasm with unmodified asparagines in NxS/T consensus sequences. The unedited NRF1 peptides that contain NxS/T glycosylation sites with unmodified N and not edited to D may signal that cellular N-glycosylation in a virally infected cells is inefficient because the cell is experiencing ER stress during the viral infection. HLA presentation of peptides bearing NxS/T glycosylation sites with an N at position 3 rather than the normally edited D at position 3 may indicate to the immune system that a viral infection is taking place in that cell. In this sense, the N to D edited NRF1 peptides are “healthy self” whereas the genomically encoded N peptides at NxS/T sites that are not N-glycosylated may be signs of self under viral stress. In addition, the “N not edited to D” peptides might be recognized as foreign, because among the set of self-peptides produced by unstressed cells during clonal deletion would be 12**D**456789 derived from NRF1 peptides which would then select for deletion of B cells that produce antibodies that recognize that peptide. But the 12**N**456789 peptides derived from NRF1 produced during any viral infection would not be normally synthesized and thus would be foreign and induce immune reactivity to the infected cell only. In this way, an unedited peptide would be a viral molecular pattern in analogy to the microbe associated molecular patterns (MAMP).

There have been earlier reports of N-glycosylated asparagines in viral or host proteins that are edited to aspartate and presented much more effectively than Asn by the mammalian immune system. For example, mammalian tyrosinase which mediates melanin production in the eye and melanocytes, is N-glycosylated and deglycosylated and only the deglycosylated edited peptide at the NxS/T at position 369 is presented by HLA (Ostankovitch et al, 2009). Tyrosinase (369)(NGT) was not presented on HLA because unglycosylated tyrosinase is degraded rapidly, but the N-glycosylation followed by deglycosylation and editing altered the selectivity of tyrosinase processing by the proteasome, enhancing the production or survival of Tyrosinase 369 DGT. This deglycosylation and editing depends on NGLY1 deglycosylation activity (Altrich-VanLith et al, 2006) The N to D edited peptide of tyrosinase is abundant in the HLA-bound proteome of human cells reported by the Monash group (Mei et al, 2020).

## The frequency of N to D substitutions in *C. elegans* N-glycosylated proteins vs other proteins

From the linguistic structure of the genetic code, asparagine codons **A**AU/C are a single nucleotide transition mutation away from the aspartic acid codons of **G**AU/C. Asn and Asp residues also have similar physical properties, often located at protein surface, and their mutual substitution usually do not have deleterious effect on protein folding or function: N to D amino acid substitutions are relatively frequent in PAM250 or BLOSUM62 matrices based on the statistics of residue substitutions among protein homologues. In those substitution matrices, N to D, S, and H substitutions are most common and roughly equally frequent across aligned protein sequences. The codons for S (A**G**U/C) and H (**C**AU/C), are also a single nucleotide difference from N (AAU/C). If the protein editing of some N-glycosylated sites of proteins to D by NGLY1/PNG-1 N-glycanase has selective advantage such that in a distinct species a genomic N to D substitution might also be advantageous, then the frequency of N to D mutations in phylogeny will be increased in the subset of N’s located in the context of NxS/T glycosylation sites. We therefore searched in three datasets, endoplasmic reticulum annotated proteins, which are most likely to be N-glycosylated, nuclear proteins, which are unlikely to be ER-transiting and N-glycosylated, and coronavirus proteins, many of which transit the ER, for the frequency of N to D substitutions in phylogeny in NxS/T N-glycosylation sites compared to conserved aligned N’s. Analyzing the patterns of amino acid substitutions in the multiple sequence alignments of 414 ER proteins from *C. elegans* and their homologues among divergent nematode species, we found that the frequency of N to D substitutions at NxS/T sites was 1.7 times higher than the frequency of these substitutions across all Ns (Figure 2C). As a comparison, in the multiple sequence alignments of 2900 nuclear proteins from *C. elegans* and their homologues, the frequency of N to D substitutions at NxS/T sites (11%) was slightly lower than the frequency of D substitution across all Ns (12%). Thus, this analysis favors the model that a cycle of ER localization, N-glycosylation of NxS/T sites in the ER, followed by deglycosylation of those N-glycosylated sites by NGLY1/PNG-1 can be replaced by an aspartic acid substitution mutation at those NxS/T sites (to a DxS/T) in other nematode species. While the N to D protein edit via N-glycosylation and deglycosylation depends on a complex pathway of N-glycosylation followed by deglycosylation in different cellular compartments, the genomic change to a D at that position bypasses the need for N-glycosylation or NGLY1/PNG-1 deglycosylation. NGLY1/PNG-1 converts asparagine to aspartic acid in a reaction that transfers the oxygen from water to the amino acid side chain to release the amide linked to the N-glycan carbohydrate that is hydrolyzed in the deglycosylation. The oxygen of water becomes the oxygen of the new aspartic acid sidechain carboxylate. Thus any species that diverges to an ecological niche with exposure to dessication, a challenge to many organisms (and common in bats), may escape the need for efficient NGLY1-protein editing with an NxS/T to DxS/T mutation.

## The viral resistance of humans and mammalian cell lines deficient in NGLY1/PNG-1 deglycosylation function

Human mutations in the NGLY1 N-glycanase deglycosylation enzyme have been known for almost a decade (Need et al, 2012). There are about 60 patients with NGLY1 deficiency who are wheelchair-bound with multiple neural deficits. Some of the molecular lesions in NGLY1 are null alleles and others are reduction of function alleles; the patients with heterozygous stop codons define the NGLY1 null phenotype. The patients show developmental delay, hypotonia, seizure, peripheral neuropathy, often no tear production or alacrima. The molecular identification of NGLY1 as an N-deglycosylation defect suggested that the defective degradation of very abundant N-glycosylated proteins that emerge from the ER or Golgi was the underlying molecular pathway that causes the disease symptoms. But the *C. elegans* genetics and the cancer dependency correlation of NGLY1 with SKN-1A strongly supports a role of NGLY1 in proteasomal homeostasis.

There are hints from the NGLY1 patients that deglycosylation and N to D protein editing by NGLY1 is salient to viral life cycles by reports of viral immunity in the NGLY1 patients and in NGLY1 deficient cell lines. The parents of these children also report profound resistance to the common colds and flu in their NGLY1 homozygous children (Lam et al, 2017). Consistent with NGLY1 function in viral life cycles, an RNAi screen for human gene inactivations that confer cell culture immunity to Enterovirus 71, an RNA virus, identified NGLY1 as a gene activity needed for viral replication (Wu et al, 2016). Support for the reports from parents of viral resistance in the NGLY1 children emerged from the comprehensive characterization of 12 of these patients which showed aberrantly high antibody titers toward rubella and/or rubeola following standard childhood vaccination (Lam et al, 2017). One explanation is that defects in the normal NGLY1 N to D editing of viral proteins, including Rubella, causes aberrant presentation of viral antigens or aberrant selection of antibodies and cytotoxic immune cells. The unexpected increase in measles vaccination antibody titres of the NGLY1 children suggests that either the defect in viral life cycles somehow intensifies immune response to viruses or that the failure to increase proteasome abundance during an immunization somehow intensifies antibody response. Titres of antibodies after an immunization depend on proteasome processing and presentation of viral peptides by the HLA complex; disruption of the proteasome response could affect processing of antigen peptides and the literature already shows that N to D editing of processed peptides favors HLA antigen presentation of viral peptides. But the NGLY1 patients are predicted to be deficient in NGLY1-mediated N to D editing, suggesting that their HLA presentation of N to D edited antigen peptides may be decreased relative to normal. Because NGLY1 is expected to mediate the editing of human or viral proteins that are presented by HLA (Mei et al, 2020) and because of the clear involvement of HLA-presented protein fragments in B cell immune selection, a simple hypothesis is that protein deglycosylation and N to D editing is deeply involved in immune function as well.

The deglycosylation of glycosylated proteins was traditionally considered a prelude to their proteasomal degradation. But for immune function, processing to peptides is a key step in immune presentation of host and viral peptides. And many of the proteins secreted by cells are involved in antibacterial and antiviral immune function. So the N-glycosylation may well be the battlefield between viruses and the immune system, with measures, countermeasures, countercountermeasures ad infinitum. With the viral immunity reports of the NGLY1 patients and the finding that so many viral proteins appear to use N to D editing of glycosylated N’s, a very different view of N-glycosylation and deglycosylation is emerging. Instead of a pathway towards simple proteasome destruction of missorted glycosylated proteins, now it can be viewed as a biological process of protein editing. And given that there is a mechanism that edits N-glycosylated asparagines to aspartate, biology and evolution have exploited it for the past billion years of eukaryotic evolution.

The high level of antibodies to measles after vaccination of NGLY1 homozygous children also suggests that this virus may use N to D editing in its battle with the host immune system. Antibodies themselves are N-glycosylated and may be edited N to D as they are secreted. Antibody genes are rearranged during maturation and very highly expressed, which could cause proteasomal stress akin to aneuploidy. Many of the cell surface molecules that mediate adhesion between immune signaling T helper cells and B cells that produce humoral immunity are also N-glycosylated and may be edited N to D by NGLY1. This N to D editing of antibodies and immune cell adhesion factors may be defective in NGLY1 patients. In addition, unlike NGLY1 competent children, the set of self peptides to which their immune system defines as self in immune tolerance in the NGLY1 homozygous patients will not include N to D edited peptides of N-glycosylated proteins. In addition, the glycosylation state of their many secreted, N-glycosylated proteins may be enhanced, possibly enhancing viral recognition of glycosylated receptors.

## Phylogenetic evidence for N to D editing in SARS-CoV-2

Just as the pattern of N to D substitutions in NxS/T N-glycosylation sites in phylogenetic comparisons of SKN-1A between nematodes and NRF1 between vertebrates could predict such edits in the natural function of these proteins, we searched for N to D substitutions in the SARS-CoV-2 coronavirus compared to other coronaviruses. In SARS-CoV-2 coronaviruses, the S glycoproteins are N-glycosylated at 22 sites, many of which can be seen in the cryo-EM map (Walls et al, 2020). Other viruses, including HIV-1, influenza, Lassa, and Ebola N-glycosylate envelope proteins with host-derived glycans to prevent antibody recognition of the underlying protein surface. The importance of N-linked glycosylation of envelope proteins in immune evasion has been observed across many viruses including Hepatitis C.

We surveyed NxS/T or DxS/T sites among the 29 open reading frames of SARS-Cov2 analyzed the patterns of their amino acid substitutions among close protein homologues in related coronavirus strains and species (Table 2). When we applied this analysis to SARS-Cov2 NxS/T sites comparing them to dozens to hundreds of close homologues detected in other coronaviruses (some protein regions are more broadly conserved across viruses than others), we detected 57 N-glycosylation or D-substituted N-glycosylation sites in SARS-CoV-2 proteins that are candidates for N to D editing by NGLY1 during infection by the coronaviruses (Table 2). The Spike protein sites showing N to D substitutions in various coronaviruses constituted 13 of these sites but multiple RNA dependent RNA polymerase and polymerase cofactor sites were also detected (Table 2). We also annotate in Table 2 the propensity for N to D substitution at individual NxS/T site or D to N substitution at individual DxS/T sites among close homologues of SARS-Cov2 proteins in other betacoronavirus genomes (compared in the percent D and percent N columns). A particularly striking example is the glycosylation site at N657 of the Spike protein of SARS-Cov2, where the D substitution frequently occurs in various coronaviruses from bats and only N or D is found at this position (Figure 3A; Table 2).

**Figure 3.**
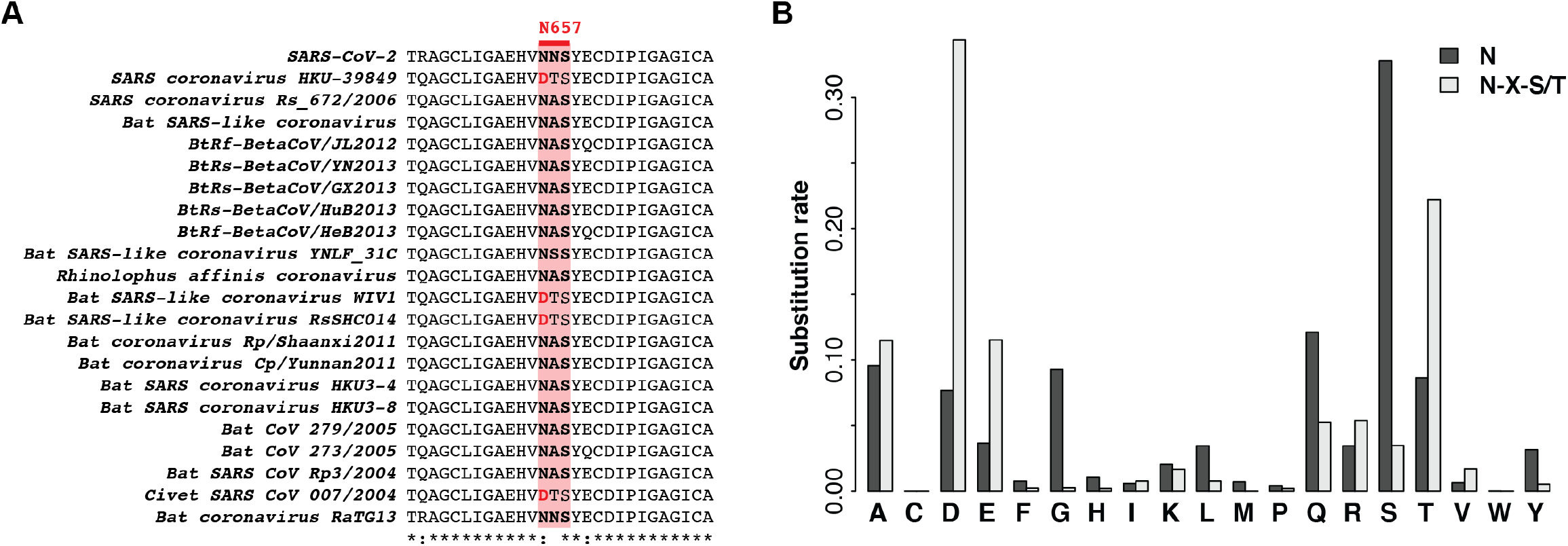
Among coronavirus homologues of SARS-Cov2 spike protein, asparagine to aspartate substitutions are strongly enriched at N-glycosylation sites. A. An example of multiple sequence alignment in the vicinity of N-glycosylation site in the spike protein of SARS-Cov2, with an increased rate of substitution to D among related coronaviruses from bats. B. Among all coronavirus homologs SARS-Cov2 spike protein, the substitution rate of N to D residue is significantly increased at glycosylation motifs compared to overall N to D substitution rate in these proteins. The bar plot shows substitution rates of N to other residues for glycosylation motifs and for all N residues.

Analyzing the full-length multiple sequence alignment of coronavirus Spike proteins, we found that the sites with NxS/T glycosylation consensus motif had a 3 times higher N to D substitution rate than the average rate for all N sites in these proteins (Figure 3B). The coronaviruses of bats show a high frequency of these N to D substitions; the diet, high metabolic rate, and possible dehydration of bats may inhibit steps in N-glycosylation that depends on N-dolichol lipid synthesis or the water hydrolysis step in NGLY1 deamidation to select for viral genomic N to D substitutions.

To further validate this method, we analyzed all NxS/T sites in the *C. elegans* proteome and assessed their N/D substitution rates in multiple sequence alignments of close homologues among 10 different *Caenorhabditis* strains (Supplementary Table 1). For SKN-1A, we could easily detect the NVS to DVS substitution between *C. elegans* and *C. brenneri* or *C. tropicalis*. In addition, we could detect patterns of N to D substitution in NxS/T glycosylation sites of *C. elegans* antiviral Argonaute proteins NRDE-3, WAGO-1, RDE-1, PRG-1, SAGO-2, WAGO-10, as well as the RNAi factors DCR-1, MUT-7, and PGL-1 (Supplementary Table 1). The N-glycosylation of the Argonaute protein is consistent with the original purification of mouse Argonaute, the founding member of the family that presents miRNAs to target mRNAs as an ER-localized protein (Cikulak et al. 1999). Similarly, in Arabidopsis, Argonaute localizes to the ER to regulate translation of mRNAs that encode secreted proteins (Li, et al, 2013).

A hint for the function of N to D editing of viral proteins emerged from the discovery of N to D editing of glycosylated asparagines of the E1 epitope of the Hepatitis C virus glycoprotein: a peptide epitope EG**NAS**RCWVA from the genomic sequence of the virus is 1000x less potent at inducing antiviral cytotoxic T cell immunity compared to a mutant where the asparagine in the NAS glycosylation site was mutated to aspartic acid, EG**DAS**RCWVA, raising the potency of this peptide by 1000x. Similarly, if the cells were treated with the N-glycosylation inhibitor tunicamycin, the potency of the EG**NAS**RCWVA peptide for immune rejection of the viral infection was strongly enhanced, showing that the N-glycosylated peptide evades immune recognition (Selby et al 1999). Thus, deglycosylation may be an immune adaptation to the viral strategy of N-glycosylation of envelope proteins to evade immune surveillance.

E1 epitope of the Hepatitis C virus glycoprotein

232-EG**NAS**RCWVA-241

232-EG**D**ASRCWVA-241

## Evidence for N to D editing in the public epitopes of viruses that are recognized by antibodies in the Elledge lab Virscan

Antibody immunity is a key step in viral immunization. Steve Elledge’s lab has explored the viral epitopes from a wide collection of viral genomes to detect antibodies in serum samples, in some cases life-long, that surveil for these viruses decades after an infection or immunization. In a comprehensive screen of potential viral 56mer peptides inferred from the genomes of 206 species of human DNA or RNA viruses screened individually as peptides displayed on a bacteriophage coat protein, the Elledge lab explored the viral infection history of 569 diverse human subjects (Xu, et al, 2015). From the patterns of 56mer test peptides that bound to antibodies in microliter blood samples from patient samples, they viewed the viral infection/immunization history of these patients and the detailed viral peptides targeted by antibodies. They found many examples of multiple independent immune responses in individuals were antibodies that bind to the same immunodominant peptide from a particular virus. These public epitopes could reflect peptide sequence motifs that rearranged antibody genes prefer to bind, and thus may trigger B cell expansion if a high affinity antibody rearrangement and hypermutation occurs. Alternatively, and more fiendish, these public epitopes could reflect an evolutionarily advantageous viral diversion tactic to trick the host immune system into an impotent antibody response that saturates the ability to mount a more effective immune response. That is, the peptide produced by the virus could be a form of propaganda rather than real information about protein domains needed for virulence that the host immune system should target.

We searched through the Virscan peptides to test by phylogenetic analysis if these antigenic peptides show indirect evidence of N to D editing in the pattern of N to D substitutions in the phylogeny of related viruses. We inspected the sequences of these public epitopes among the coronaviruses first (Quiat et al, 2020). Out of 408 epitopes identified from dozens of diverse DNA and RNA viruses, 23 were from coronaviruses (shown in Supplemental Table 2, data from Xu et al 2015 and Quiat et al, 2020). We compared the 56 mer peptides across the betacoronaviruses to search for N to D variation in NxS/T possible glycosylation sites. One of the 56mer peptides that human antibodies bind is shown below:

1. Bat coronavirus 1B ORF1ab RNA dependent RNA polymerase

CIAARDVVVTNL**NKS**AGYPLNKFGKAGLYYEALSYEEQDALYAVTKRNILPTMTQL

This bat coronavirus peptide derives from the RNA-dependent RNA polymerase that is processed from ORF1ab of the coronavirus. The N in the NKS glycosylation site of this bat coronavirus 1B RdRp has mutated to a DKS in the SARS-CoV-2 coronavirus (Figure 5A, shown in red). The immunodominant epitope was mapped with overlapping peptides by the Elledge group to the 15aa peptide sequence NL**NKS**AGYPLNKFGK in the bat coronavirus.

2. Human coronavirus NL63 Replicase polyprotein 1ab (pp1ab) (ORF1ab polyprotein)

A second public epitope from Human coronavirus NL63 derives from the homologous region of the RdRP in human NL63, and shows the same N to D edit in SARS-CoV-2 and many other betacoronaviruses (Figure 5B, Supplemental Table 2). This sequence is mutated in SARS-CoV-2 to a DKS (Figure 5B, shown in red). Based on antibody recognition of overlapping 56mers, the public epitope of this 56 mer Human coronavirus NL63 56mer is **NKS**AGWPLNKFGKAS, highly homologous to the Bat coronavirus public epitope above. Thus, two independent coronaviruses, Human NL63 and Bat coronavirus 1B bear an NKS motif that is highly immunogenic and mutated to DKS in SARS-CoV-2. The N to D changes of this site in coronavirus evolution predict that the NKS sequence of this peptide is N-glycosylated in the ER, and that the glycosylated asparagine at the NKS is deglycosylated in the cytoplasm by NGLY1, removing an amine in the process. It is possible that the actual epitope presented to the bat immune system by HLA is the deglycosylated protein edited **DKS**AGWPLNKFGKAS peptide of NL63, whereas in SARS-COV2, no NGLY1 editing is required because the genome sequence is mutated to DKS. Supplemental Figure 2 shows the phylogenetic variation of that SARS-CoV-2 DKS to NKS of this ORF1ab RdRP protein in other coronaviruses. Note the many N to D substitutions.

## Mass spectroscopy endorsement of SARS-CoV-2 deglycosylations

While the set of protein modifications that SARS-CoV-2 proteins undergo have not been as exhaustively analyzed by mass spectroscopy as the human proteome for N to D edits, predicted N-glycosylation modifications in coronavirus spike proteins have been observed by mass spectroscopy. And there is genetic support for the editing of the N-glycosylated asparagine to aspartic acid at some positions during deglycosylation, allowing a mutant viral spike protein in the related chicken coronavirus infectious bronchitis virus (IBV) to be functional; D to N substitution mutations at 13 out of 14 of the 27 validated N-glycosylation sites tested in the IBV coronavirus Spike protein were competent for both viral replication and induction of cell fusion (Zheng et al 2018). We explored how the candidate glycosylation sites of SARS-CoV-2 correlated with immune epitopes at the Immune Epitope Database (IEDB) https://www.iedb.org/home_v3.php).

A predicted N to D variation from SARS-CoV-2 is the spike protein NxS/T site at 657 that mutates to DTS in SARS CoV.

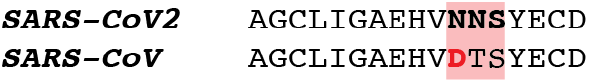

This peptide derives from the E2 glycoprotein precursor (633-649) of SARS-CoV-2 coronavirus Tor2 which is positive in ELISA assay for antibody binding after immunization of rabbit and human infection in IEDB. Another region of SARS-CoV-2 ORF1ab peptidase and shows the N to D variation in Table 2 detects a peptide in IEDB in Coronavirus is YLDGA**DVT**KIKPH from the original SARS-CoV that binds to HLA-DRB1*01:01 with an affinity of 2 nM. The region flanking this site from SARS-CoV-2 is DMSMTYGQQFGPTYLDGA**DVT**KIKPHNSHEGKTFYVLPNDDTLRVEAFEYYHTTD

Multiple peptide epitopes from SARS-CoV bind HLA with nM affinities from IEDB data.

LRSEAFEYY,NTNLHTQLVDMSMTY,PTYLDGADVTKIKPH,SEAFEYYHTL,TQLVDMSMTY

Another sequence from SARS-CoV-2 showing N to D variation between coronaviruses maps to the RdRP and the variation is shown in the Figure above. 4891 IVNNL**DKS**AGFPF from SARS-CoV-2 RdRp

Retreiving the region flanking the DKS at 4891 and comparing to Bat coronavirus 1B RdRP shows a pattern of D to N variation. The D in the SARS-CoV-2 coronavirus RdRP is changed to an N in the NxS/T glycosylation site of this bat coronavirus 1B RdRp. The comparison below bolds the DKS to NKS variation.

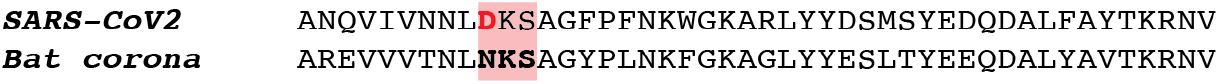

A search with a region flanking the predicted N to D variation in the SARS-CoV-2 RdRp detects multiple peptides from the IEDB that bind to HLA (RLYYDSMSY,SYEDQDALF,DQDALFAYTK,GGCINANQVIVNNLD, IPTITQMNLKYAISA,YEDQDALFAY,LFAYTKRNVIPTITQ)

Another example, the RQHLK**DGT**CGLVE predicted peptide of the SARS-CoV-2 NSP1 replicase subunit retrieves a peptide with a D to N substitution this is a bona fide HLA binding epitopes from SARS-CoV (the original SARS):

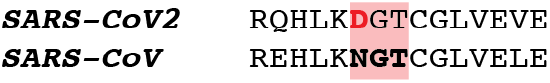

The peptide epitope from the Replicase polyprotein 1ab from Severe acute respiratory syndrome-related coronavirus (Human coronavirus (strain SARS))., tested in on HLA-DRB1*01:01 which binds at 7.2 nM.

## Other Virscan RNA viruses showing evidence of N to D editing

There were more examples of public epitopes to viruses unrelated to the coronaviruses that showed NxS/T N-glycosylation site changes to DxS/T in related viruses. We annotated NxS/T sites in the 211 unique 56mers that bound to human antibodies among the entire proteomes of the wide range of viruses analysed in Virscan. We found 47 NxS/T sites in these peptides, and searched in related viruses for examples of N to D substitutions. The sites with the most D substitutions were from the Human rhinovirus B70 polyprotein, influenza virus hemagglutinin, human papillomavirus type 3 major capsid protein L1, Hepatitis C virus polyprotein, and HIV envelope glycoprotein.

1. Rhinovirus B in the viral coat protein VP2 (common antibodies to his antigen)

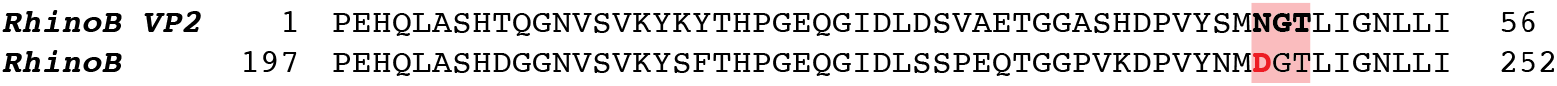
2. A peptide from influenza virus that antibodies from many individuals bind to also shows a high frequency of N to D substitutions in other related viruses is the hemagglutinin protein, a hot spot for viral pandemic variation. Many humans tested have antibodies to this epitope (Quiat et al, 2020); we all catch the flu. Hemagglutinin Influenza A virus (strain /Memphis/3/1988 H3N2)

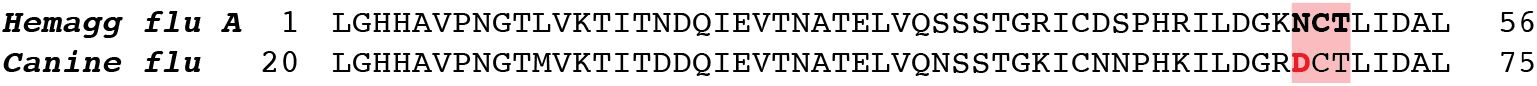
3. Human immunodeficiency virus 1 Envelope glycoprotein gp160 is shown below with potential NxS/T sites bolded:

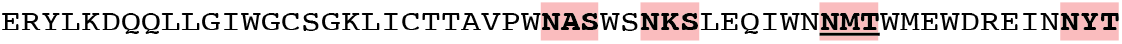

The NMT (underlined) site has been detected as N to D deamidated in a cell culture assay of HIV HLA peptides produced. The IEDB reports a deamidated peptide KSLEQIWN**N**MTW Deamidated (N)@9 thus an N to D edit in vivo and bound to HLA B*57:01 Human immunodeficiency virus 1 cellular MHC/direct/fluorescence qualitative binding Positive (Ramarathinam et al, 2018).

The viral gene peptides containing NxS/T sites that were tested by antibody immunoprecipitation of viral epitopes expressed in bacterial phage fusion proteins in Virscan are not N-glycosylated in the experiment because these mammalian viral epitopes are synthesized by E. coli as it produces the fusion proteins of the T7 phage (Xu et al, 2015). E. coli does not have an STT3 dolichol transferase, a eukaryotic specialization for N-glycosylation. So that the human antibodies from sera that react for example with the **NKS**AGWPLNKFGKAS epitope of coronavirus NL63 will not be inhibited by N-glycosylation of the peptide which might obscure that epitope if it was produced in the eukaryotic cell. But it is possible that the human antibody might actually target the NGLY1-edited peptide **DKS**AGWPLNKFGKAS, which might have been the actual immunogenic peptide produced during the human viral infection and displayed on HLA in immune cell selection. The DKS peptide produced in vivo after N-glycosylation and deglycosylation may have been the original potent immunogen for the production of long lived antibodies that recognize the unglycosylated bacterially produced **NKS**AGWPLNKFGKAS epitope. Interestingly, Virscan has tested many phylogenetically related viruses, some of which show N to D substitutions of NxS/T sequences in one viral genome. Thus Virscan does compare N peptides that are not N-glycosylated in E. coli and D peptides from related viruses that might mimic deglycosylated and deamidated N-glycosylated peptides from a related virus.

## N-glycosylation and deglycosylation as targets for antiviral therapies

If N to D editing of the many viral proteins that are N-glycosylated in the ER is required in viral life cycles, a simple prediction would be that drugs that inhibit either N-glycosylation or deglycosylation by NGLY1 might inhibit viral infections. Loss of N-asparagine deglycosylation by a *png-1* (the NGLY1 orthologue) null mutation in *C. elegans* is viable and quite healthy, unless proteasome function is compromised by a proteasomal reduction of function mutation or by a drug that inhibits proteasomal function. Thus the therpeutic window between efficacy and toxicity would be wide in *C. elegans*. But loss of N to D editing via a mutation in NGLY1 in mice is lethal, and the multiple disabilities in the NGLY1 children show that it is not benign to lose NGLY1 deglycosylation activity. And while NGLY1 human patients show viral resistance, many organ systems are compromised by the human NGLY1 null mutation (Lam et al, 2017) strongly suggests that any drugs that inhibit NGLY1 could have toxicity. But partial drug inhibition of NGLY1 activity may find a sweet spot between its essential developmental function and its role in viral life cycles. There is a drug that inhibits NGLY1, the thiol reactive caspase inhibitor z-FAD (Tomlin et al, 2017), but it has not been used on humans or virally infected animals. In cell culture, z-FAD caused resistance to the positive strand RNA virus enterovirus 71 replication (Wu et al, 2016). NGLY1 inhibitors such as z-FAD (Tomlin et al, 2017; Wu et al, 2016) delivered to the airways may show less toxicity. In addition, siRNA inhibition of NGLY1 only in the accessible lung tissue could be a therapeutic avenue.

One attractive theory for the selection for N to D substitutions in animal or viral evolution that was not endorsed by comparative genomics is that D substitutions could be selected in hosts that are deficient in NGLY1 deglycosylation. The chiropteran bats and pangolin, suspected hosts of SARS-CoV-2 viruses which show much N to D variation, all encode NGLY1 deglycosylation and STT3 N-glycosylase homologues. Interestingly however, while NGLY1/PNG-1 is conserved in nearly all animals, plants and fungi, it has been lost in many basidomycota, most protists, and a few parasitic nematodes (Tzelepis 2017; Sadreyev 2015)(Figure 4).

**Figure 4.**
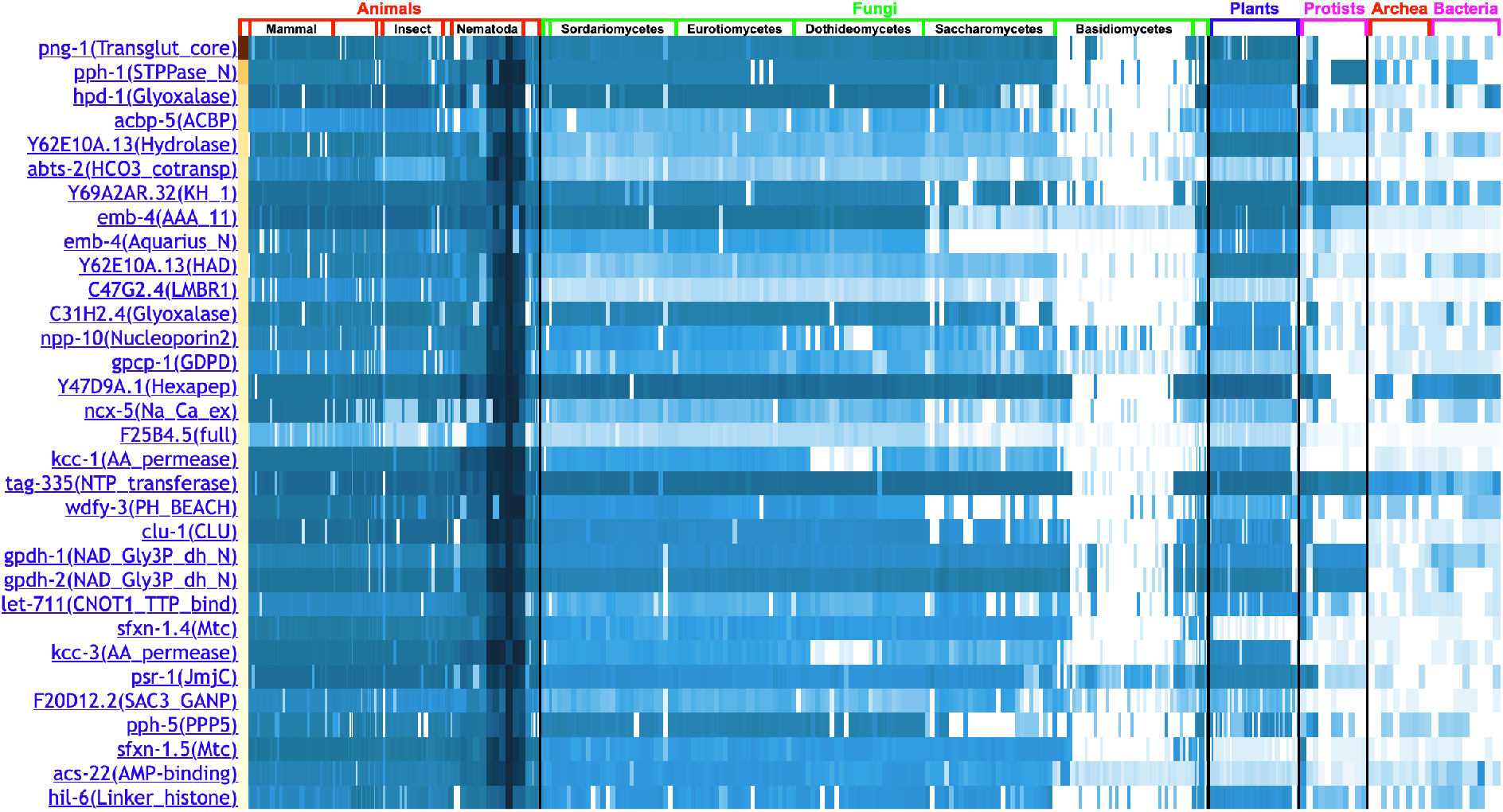
Heatmap of conservation patterns of while NGLY1/PNG-1 proteins across phylogenetic clades. Close NGLY1/PNG-1 homologs, shown in dark blue, are present across animals, plants and fungi, but not in most basidomycota and protists. They were also lost in a few parasitic nematodes. Shown on this phylogenetic profile are other *C. elegans* protein domains with no homology to NGLY1/PNG-1 that show a similar pattern of loss in basidomycota and protists; such correlated losses can reveal other genes in the deglycosylation pathway or pathways that intersect with NGLY1/PNG-1 deglycosylation and protein editing (Tabach et al, 2013).

**Figure 5.**
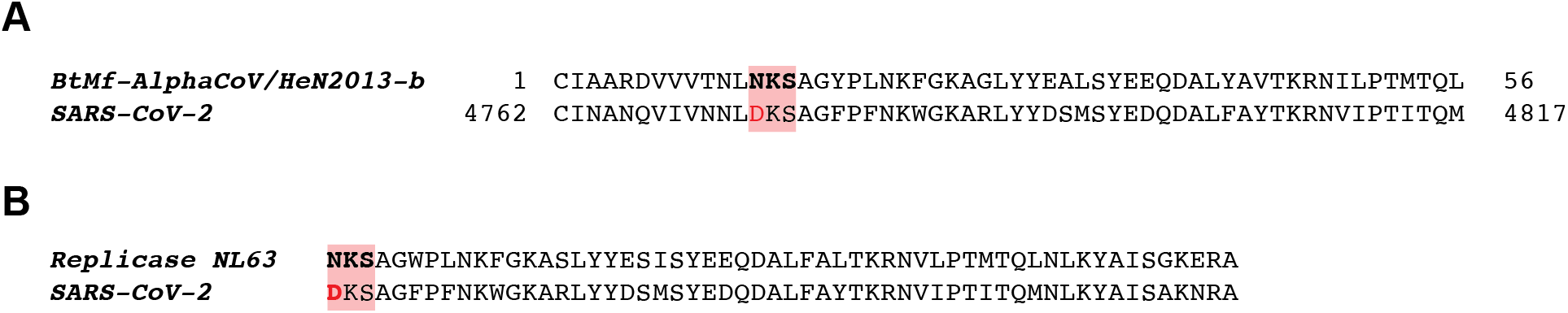
Public coronaviral epitopes commonly recognized by human antibodies are often subject to N/D substitution at N-glycosylation motifs. Public epitopes of coronaviruses with N to D substitutions at glycosylation motifs among close homologues. A-C: Coronavirus epitopes. A. Epitope of RNA dependent RNA polymerase (ORF1ab) from bat coronavirus 1B. B. Epitope of RdRP from human coronavirus NL63 shows N to D substitution in Covid 19 virus.

In addition, the PNG1 deglycosylation enzyme of many but not all Ascomycota are missing the predicted catalytic residues for the deamidation of client N-glycosylated proteins (Maerz 2010), but have clearly conserved the N-glycosylation binding activity of PNG1. For example, the Neurospora crassa PNG1 missing the catalytic triad is no longer is capable of deamidation, but is required for normal conidial growth because mutations in the conserved but catalytically inactive N-glycosylation binding domain show conidial defects (Maerz 2010). PNG1 of Neurospora may bind to protein glycans at the fungal cell wall to mediate conidial growth. In contrast, the budding yeast S. cerevisiae has a fully catalytic PNG1 which is predicted to deamidate and edit N to D in an unknown client N-glycosylated protein. The S. cerevisiae client proteins have not been identified but may be analogous to the ER-transiting SKN-1A/NRF1 transcription factor proteins.

N-glycosylation is used by thousands of host proteins; loss of STT3 gene function in N-glycosylation, the enzyme that actually transfers glycosyl groups from n-dolichol to asparagine residues on client N-glycosylated proteins, is lethal in many animals, including *C. elegans*. Tunicamycin inhibits eukaryotic N-glycosylation by STT3-mediated N-glycosylation and is toxic to eukaryotic cells. Thus, inhibition of N-glycosylation is not an attractive candidate for an anti-viral therapy. Low doses of tunicamycin or aerosol tunicamycin could inhibit Covid viral life cycle in the lung without as much toxicity. Statins also inhibit N-glycosylation because they inhibit HMG CoA reductase, a key step in the synthesis of mevalonate, a precursor to N-dolichol in the N-glycosylation pathway, as well as cholesterol in that biosynthetic pathway. Statins have emerged from screens for approved drugs that inhibit the replication of hepatitis C virus, an RNA virus (Kim et al, 2007; Shrivastava-Ranjan et al, 2018). But statins are absorbed in the intestine and immediately transported to the liver, the center of cholesterol synthesis and regulation where they inhibit the production of cholesterol more than the other outputs of the mevalonate pathway. But statins inhibit the N-glycosylation of the insulin-like growth factor I and II receptors in the placenta (Forbes et al, 2015), and a major side effect of statins is muscle pain probably caused by inhibition of coenzyme Q synthesis, suggesting that the drug inhibits more than only the cholesterol output of the mevalonate pathway and in cell types outside of the liver. Hundreds of millions of people take statins to lower their cholesterol. However, our analysis of how patients treated with statins fare in SARS-Cov-2 infections showed an increased mortality and morbidity, suggesting that inhibition of N-glycosylation is not a promising avenue for viral resistance (Ravi Thandhani, Yuchiao Chang, Gary Ruvkun, manuscript in preparation).

## Do viruses need N to D protein editing by NGLY1/PNG-1?

A model based on our viral phylogenetic evidence of N to D editing (for example at N657 of the SARS-CoV-2 Spike protein or a similar N to D pattern at D4891 of Orf1ab of SARS-CoV-2 and related coronaviruses which encodes the RNA dependent RNA polymerase protein) is that NGLY1-mediated protein edits are necessary for efficient production and shedding of viral particles and these edits do not occur in the NGLY1 patients. This is also consistent with the identification of NGLY1 in an RNAi screen for mammalian gene inactivations that disrupt enterovirus life cycles (infection/membrane fusion, RNA replication, viral packaging, cell biology, or budding) (Wu et al, 2016). The resistance of NGLY1 patients to viruses could be tied to the evidence for viral N to D editing that our phylogenetic analysis of virus genome sequences supports. The N to D edits predicted by the phylogenetic analysis of the Spike protein N657D change, would be expected to not occur in these patients. In this sense, the viral life cycle may be compromised by the lack of N to D editing in these patients.

From the perspective of the virus, is N to D editing something useful for the viral life cycle so that the edited proteins actually are incorporated into the viral particle? An attractive hypothesis is that extremes in membrane torsion that are required for viral budding or viral membrane fusion during infection may require deglycosylation of some N-glycosylated Spike proteins. This hypothesis emerges from the observation that NGLY1-deficient humans show defects in tear gland and adrenal gland function, organs with highly convoluted secretory surfaces that may be defective in the absence of NGLY1-mediated deglycosylation (Lam et al, 2017; van Keulen et al. 2019). Proteomic mass spectroscopic analysis of SARS-Cov-2 viral particles may reveal if N to D editing (the addition of .98 daltons by the OH in place of the NH2 to peptides predicted from the genome sequence) of Spike protein is packaged into viruses. Similarly, proteomic analysis of any virus showing N to D variation in NxS/T N-glcosylation sites in viral phylogeny, as we pointed out above for rhinovirus B, influenza, and HIV, would demonstrate the involvement of N to D editing for viral life cycles.

But it is possible that it is the lack of N to D editing of the key known target of NGLY1, the transcription factor SKN-1A in *C. elegans* and NRF1 in mammals, that in turn activates proteasome production, is key the resistance to viruses of the NGLY1 patients. Both *C. elegans* genetics and analysis of human tumor lines suggests that the key eukaryotic cellular target for NGLY1 is SKN-1A/NRF1 to activate proteasome production. The genetic analysis of the *C. elegans* orthologue of NGLY1 showed that a key target of PNG-/NGLY1 is not the many abundant N-glycosylated proteins but instead a particular rather non-abundant ER-localized SKN-1A transcription factor which mediates regulation of proteasome homeostasis (Lehrbach et al, 2016, 2019). The human cancer genome dependency map detects a highly significant correlation between the tumor types that are dependent on NRF1, the SKN-1A orthologue, and NGLY1 (Wang et al, 2017). A simple model for the NGLY1 and NRF1 vulnerability of particular tumor cell lines is that these cell lines are aneuploid, a major challenge to proteasome as multi-protein complexes are impacted by gene copy number imbalances (Oromendia 2012). A simple model is that without NGLY1-mediated N to D modifications to the NRF1 transcription factor, the normal increase in proteasome production that accompanies the ER stress of viral infection (and that viruses may “expect” or require) will not occur. Less proteasome capacity can impact viral assembly and budding from cells.

## An outline of immunological experiments to seek evidence that N to D edits by NGLY1 are salient to anti-SARS-Cov-2 immune responses

The N to D protein edits on SARS-Cov-2 Spike protein for example are not known to be incorporated into viral particles. Proteomic experiments to seek N to D edited peptides would be one test of this theory, but not a functional test of a function in immune response to viral infection. N to D edited peptides are likely to be presented by HLA for immune selection of B and T cells. HLA clearly presents N to D NGLY1-edited peptides and there is good evidence that it may even prefer them (Selby et al 1999). If the N to D edited Spike protein for example is incorporated into infectious viruses or the site of viral budding, immune cell and immunoglobulin reactivity to D-edited peptides could confer more potent viral immunity.

So a key set of experiments is to challenge human sera from SARS-Cov-2-infected patients with 60-mer peptides that tile along the Spike protein, in the manner of Virscan (Xu et al, 2015), comparing peptides with the genomic NxS/T sequences to peptides carrying mutations to DxS/T. The test would ask whether 56mer peptides that are preedited in their genome sequences from encoding NxS/T to DxS/T are recognized by the antibody repertoire of an infected human. Based on the 1000x increase HLA presentation of N to D edited peptides, it is possible that during a SARS-CoV-2 viral infection, it is the NGLY1-edited N to D peptides that the immune system recognizes best (Selby et al 1999). If an ELISA or antibody binding test such as Virscan showed that a Spike protein peptide with an NxS/T substituted to DxS/T binds to sera from infected patients with a higher affinity than the genome matched peptide bearing NxS/T, it would favor that HLA binding to N to D edited viral peptides selects for the immunglobulin repertoire in an immune response. If the peptides with the D substitutions react more strongly to the sera, it is possible that the antibodies are directed against the N to D edited Spike protein HLA presented protein fragments rather than the N-glycosylated or unglycosylated NxS/T peptides that the genome encodes. More complex sorting of patient sera from those with ICU-severe SARS-CoV-2 vs asymptomatic SARS-CoV-2 infection could sort the N to D editing with distinct immune responses. Alternatively, hamster sera from virally infected hamsters, an animal model of human SARS-CoV-2 infection could be used.

In addition to testing N to D edited peptides for binding to immunoglobulin, testing them for binding to or HLA presentation on cytotoxic T cells or T helper cells would be an important test as well. Because Virscan presentation of peptides in the T7 phage capsid protein does not N-glycosylate peptides in E. coli, it may be necessary to use synthetic peptides bearing the N, D, and N-glycosylated version of each peptide. Testing for T cell responses to these peptides would be more complicated but the logic of the comparisons between the N, D, and N-glycosylated version of each peptide would be the same. Similarly, this could be tested with the rhinovirus B, influenza, and HIV virus test peptides and patient sera as well.

For SARS-CoV-2, we highlight multiple N to D substitutions betacoronavirus phylogeny that may correspond to actual NGLY1-edited viral proteins that act in antiviral immunity. The premier candidate Asn to Asp edits to make in the highly immune relevant SARS-CoV-2 Spike protein because of the high frequency of N to D substitutions across the betacoronaviruses are:

## N to D editing may be broadly important in the design of vaccines for SARS-CoV-2 as well as for most other viruses

Current immunization protocols that translate Spike protein mRNAs near the injection site may not expose the immune system to N to D edited Spike protein peptides that might be highly immunogenic in an actual SARS-CoV-2 infection and immune response. The mammalian cell-expressed Spike protein expressed out of the context of a viral infection may be N-glycosylated but the complexities of NGLY1 deglycosylation and editing may be attenuated or missing in cells that are not actively budding viral particles. A pre-edited N to D change in the genome sequence of the Spike protein may engage the immune system more like a virally-infected cell.

One test for vaccination potency would be to engineer multiple N to D substitutions into the Spike protein, for example, to then assess whether these N to D substitutions confer more potent viral immunity to hamsters for example. Immunization with an array of N to D mutated protein antigens could bypass the need for N-deglycosylation in vaccination, at least accelerating vaccine development and perhaps enabling vaccination with immunologically more potent D-edited antigens. It may enable more potent vaccination with bacterially produced peptides, which are not N-glycosylated in bacteria at NxS/T sites and not deamidated to aspartic acid in bacteria which have no NGLY1/PNG-1 homologue proteins. Vaccination with bacterially produced antigens would bypass the complexities of tissue-culture or mRNA-programmed mammalian vaccination.

More generally, if any of the 29 SARS-CoV-2 proteins is expressed in bacteria or other hosts as edited fusion proteins that are designed to change each of the 57 predicted N-glycosylation sites in Table 2 to an aspartic acid residue, 57 NxS/T to DxS/T substitutions (and of course combinations of 56, 55, etc), these engineered proteins could elicit the production of antibodies or T cells after human immunization that are more neutralizing because they select for the edited D that is a hallmark of successful expression of the SARS-CoV-2 proteins in the ER and editing to D in the cytoplasm. The same logic would apply to any NxS/T sequence in any of viral proteins from the diverse viruses discussed above.

An advantage of this approach for viral immunization is that viruses encode a dozen to a few dozen proteins, about half of which are glycosylated at countable sites, so the number of engineered changes necessary is relatively small. The NxS/T sites with N to D substitutions in related viruses can be chosen as the premier candidates for edits for immunization. However, it is also reasonable to assume that some of the highly conserved NxS/T sites are essential for the viral life cycle, for example if the N-glycosylated protein can evade immune surveillance better than any N to D substitution.

Because of the phylogenetic evidence that the NGLY1/PNG-1 editing of protein sequences has functional importance for SKN-1A/NRF1 and as outlined in this section, is likely to be used in viral life cycles as well, and because current immunization protocols do not address the probable editing and functional importance of N-glycosylated aspargines to aspartic acid in normal viral infections, we suggest that immunization with viral proteins engineered to substitute D at genomically encoded NxS/T sites of N-glycosylated viral proteins may enhance immunological response to peptide antigens. We specifically have highlighted those NxS/T sites that show a high frequency of N to D substitution in viral phylogeny as likely to be salient to viral life cycles. Because N to D edited peptides are clearly produced and presented by mammalian HLA, such peptides may more robustly activate T-cell killing or B-cell maturation to mediate more robust viral immunity. Immunization with genomically-N to D edited proteins would not require ER-localization for N-glycosylation or other cell compartment localization for NGLY1/PNG-1 N to D protein editing. Genomically substituted N to D protein vaccines could be produced in bacteria because N-glycosylation and deglycosylation which do not occur in bacteria would no longer be required to immunize with a D-substituted peptide. Thus vaccinations with bacterially-produced antigens may be efficacious.

## Acknowledgements

We thank Nic Lehrbach, Joe Goldstein, Ravi Thadhani, and Tadashi Suzuki for advice and comments that we really needed. The authors’ expertise is molecular genetics, most especially *C. elegans*, and comparative genomics. Our suggestions for the salience of NGLY1/PNG-1-mediated N to D protein editing in viral infection and immune evasion are a generalization from the discovery of NGLY1/PNG-1 protein editing of SKN-1A in *C. elegans* proteasomal homeostasis, a corner of biology not obviously connected to virology or immunology. The connection to virology was triggered by our observation of N to D substitutions in viral phylogeny that were exactly like the N to D genomic mutations in SKN-1A phylogenetic comparisons, and by the viral resistance of NGLY1 patients and the disruption of viral life cycles by NGLY1 gene inactivations in mammalian cell lines. We expected that our view of the viral life cycles through the prism of N to D protein editing was likely to be distinctive and perhaps of broader than usual interest during the Covid epidemic. We greatly appreciated the advice and comments of professional virologists or immunologists: Andrea Carfi and Kapil Bhal from Moderna, Alan Korman and Johannes Grosse from Vir, and Steve Elledge, Hidde Ploegh, Tom Rapoport, Fred Goldberg, Jack Szostak, Keith Blackwell, and Ray Chung from Harvard Medical School. We have no plans to explore the N to D editing of viral immunization ourselves, but would be excited to share any of our perspectives with virologists and immunologists who might want to ramp up such a project. We thank the Grace Science Foundation, Human Frontier Foundation, and NIH R01AG16636 for support.

## Methods

### Phylogenetic comparisons

All virus protein sequences were downloaded from NCBI: https://www.ncbi.nlm.nih.gov/labs/virus/vssi/#/virus?SeqType_s=Protein&Completeness_s=complete Redundant viral proteins were filtered using CD-hit (PMID: 23060610) with the cutoff of 95% sequence identity. To identify SARS-CoV2 homologous protein sequences among all viruses, each protein in SARS-CoV2 were split into 100 amino acid fragment sequences with 50 amino acid step, and each fragment sequence was compared against none-redundant virus sequence database using BLASTP. All identified homologous sequences were aligned using Clustal (PMID: 30976793). Protein sequences were compared by multiple sequence alignment and the frequency of asparagine changes to any other amino acid was calculated for amino acid residue positions between identified homologous virus proteins. For analysis of coronavirus sequences in regions of high conservation, for example the C terminal region of the Spike protein, 246 different viruses (mostly betacoronaviruses) could be aligned and the frequency of N to D changes were calculated between them. But for less conserved regions, for example, the N terminal region of the Spike protein, in some cases as few as 18 most closely-related betacoronavirus protein sequences could be aligned. Among those aligned protein regions, the percentage of N or D at NxS/T or DxST sites in SARS-CoV-2 was determined. For example, as shown in Table 1, for the NGT N-glycosylation site at position 282 of the Spike protein, the sequence was DGT in 16 percent and NGT in 51% of the 51 coronaviruses for which this coding region could be aligned, and the T at +3 from the N was conserved in 51% of these coronaviruses (once N is mutated to D, there is no selection for conservation of the T at +3). At Spike protein position 657 in Table 1, there are 20 alignable viral proteins from coronaviruses most closely related to SARS-CoV-2, which are N at this position in 85% of the viruses, including SARS-CoV-2, and D at this position in 15% of the alignable betacoronaviruses. To analyse the N-glcyosylation sites across the SARS-CoV-2 genome, we searched for NxS/T sites as well as DxS/T sites and then surveyed for N to D or D to N variation across other identified coronaviruses. In this way, we could infer N-glycosylation sites in other open reading frames of SARS-CoV-2, and D substitutions in related coronavirus genomes.

**Table 1.** Peptides from the NRF1 protein with N to D substitutions at N-glycosylation motifs predicted by mass spectrometry (from Mei et al, 2020).

| Peptide | Residue substitutions | Proteins Bound |
| --- | --- | --- |
| TPINQN(+.98)VSL | N to D change in cartilaginous fish. | Bound to HLA A*01:01 |
| YSAPPGDPLSTN(+.98)Y(S) | Conserved but no N to D variation in cartilaginous fish |  |
| EVN(+.98)TSASEILY |  |  |
| YLEHVGHN(+.98)HTY | N to D variation in zebrafish but not cartilaginous fish |  |

**Table 2.**
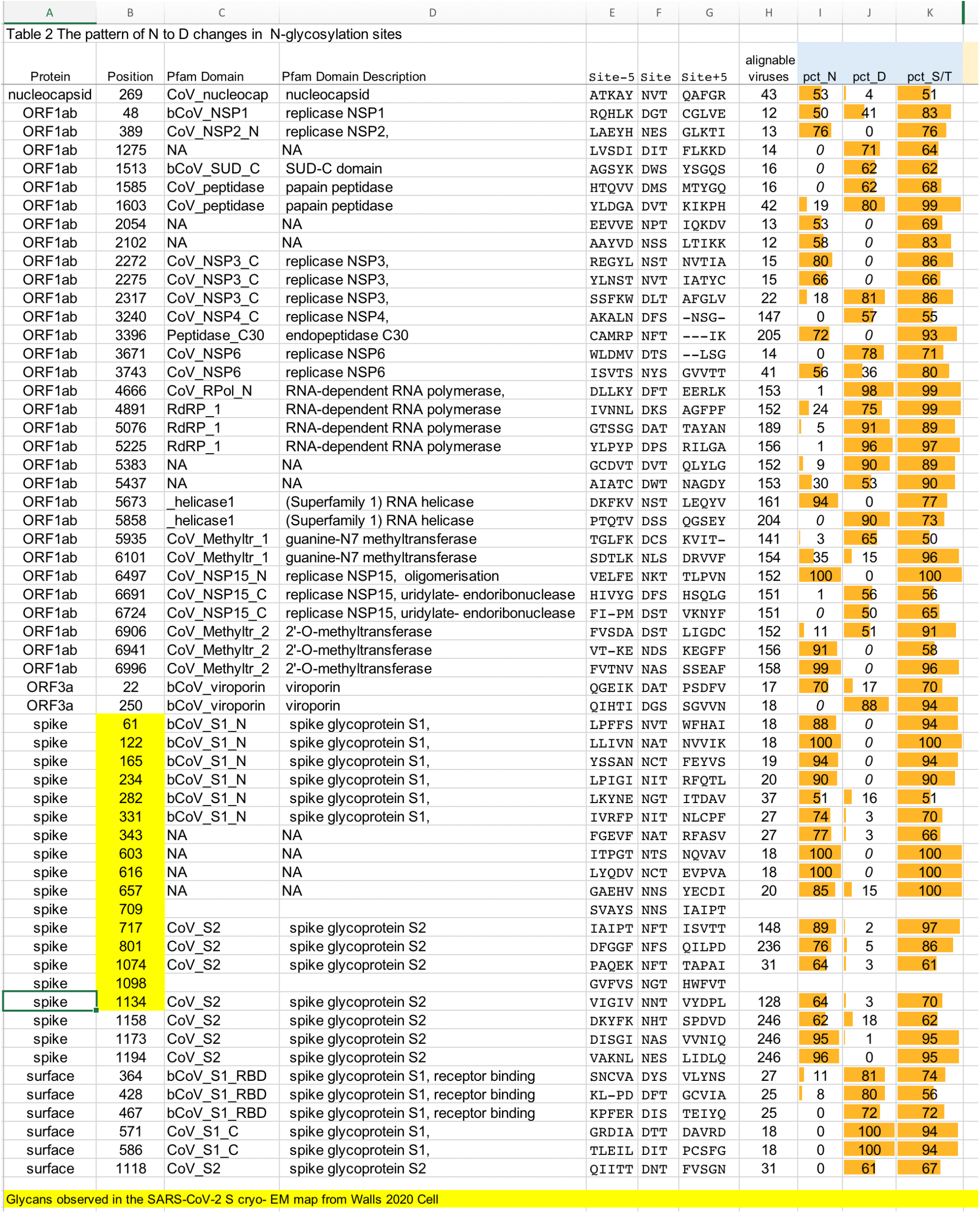
NxS/T potential N-glycosylation sites (and DxS/T sites that may be NxS/T in other coronaviruses) in SARS-Cov2 encoded proteins. Showing the 3 amino acids in the NxS/T or DxS/T site, and the number of related alignable viruses analysed, the percentage that have an N at the position or a D at the position, and the percent conservation of the S or T at +3.

**Table 3.**
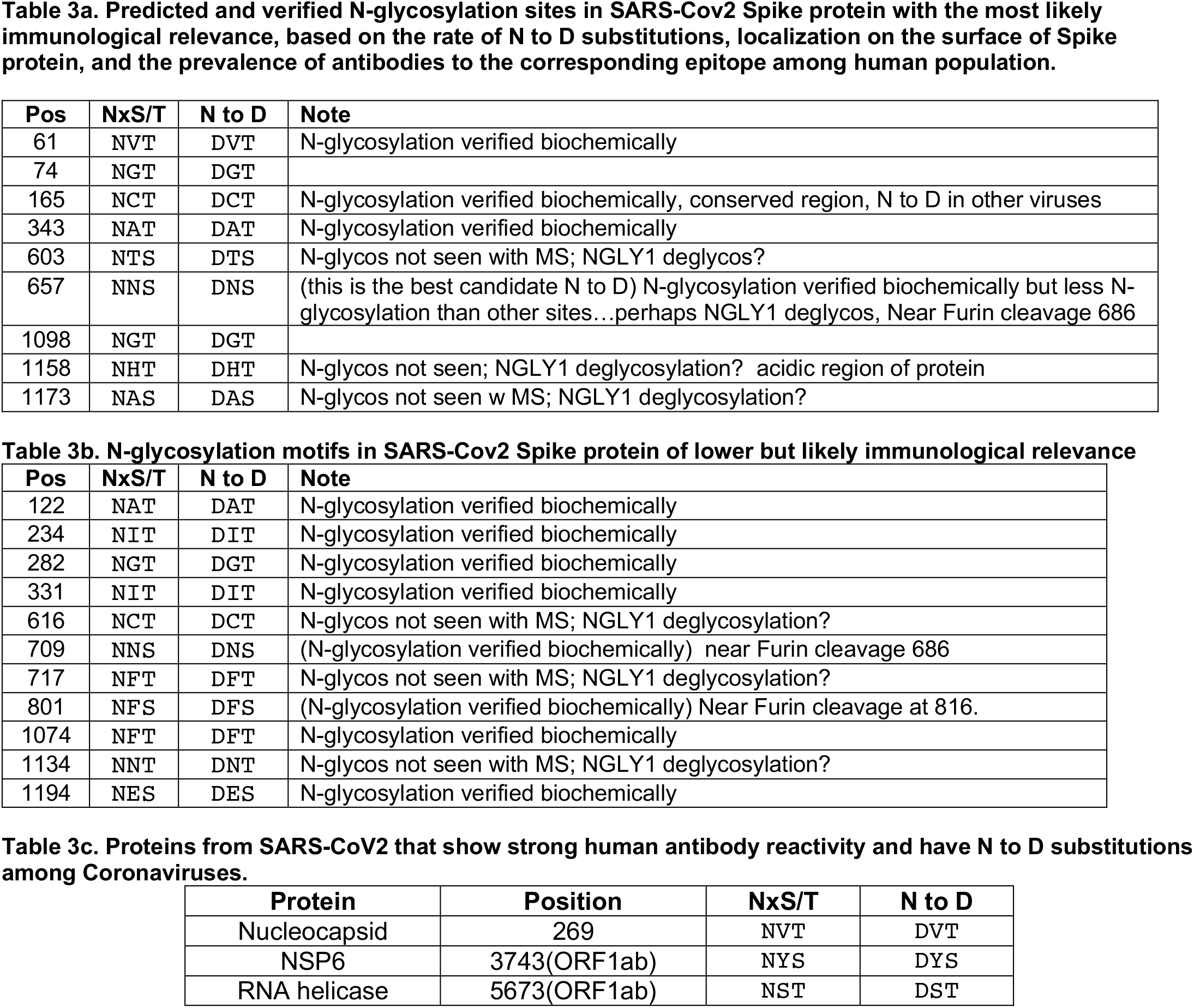
Top predicted and verified N-glycosylation sites in SARS-Cov2 Spike protein with the most likely immunological relevance, based on the rate of N to D substitutions, localization on the protein surface, and the prevalence of antibodies to the corresponding epitope among human population. Motif sequence, residue position in SARS-COV2 Spike protein, substitution sequence, and additional information about available biochemical validation data and positioning relative to the furin cleavage site are indicated.

The assessment of N-glycosylation sites in the *C. elegans* proteome involved searching for NxS/T sites in all *C. elegans* protein coding genes and comparing the conservation of those sites in 10 related Caenorhabditis species: *C. brenneri, C. briggsae, C. inopinata, C. japonica, C. latens, C. nigoni, C. remanei, C. sinica*, and *C. tropicalis*. These species are separated by up to 100 million years, based on the vertebrate fossil record-synchronized phylogenetic distances. Satisfyingly, sorting the thousands of potential NxS/T sites in the *C. elegans* proteome for those that show a high frequency of N to D substitutions in other Caenorhabditae, highlighted the NxS/T sites of SKN-1A among the top scoring N to D substitutions.

### Estimation of amino acid substitution rates

As the set of *C. elegans* ER proteins, we used 414 proteins in GO “endoplasmic reticulum” cellular component annotation (GO:0005783) from *C. elegans* (https://www.ebi.ac.uk/QuickGO). These proteins were used as queries to search for sequence homologs in the broader set of worm genomes from Wormbase (PMID: 31642470). Protein sequences of all species from Wormbase WS276 were downloaded and redundant proteins with sequence identity over 90% were filtered out using the CD-hit tool (PMID: 16731699). The resulting database was used in the BLASTP (PMID 2231712) search for the homologs of *C. elegans* ER proteins. Multiple sequence alignments of detected homologs for each C. elegans protein were constructed using CLUSTALO (PMID 21988835). These sequence alignments were used to calculate the frequency 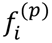 of each amino acid *i* at each position *p*, and to estimate the position-specific substitution rate 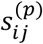 between amino acids *i* and *j* as 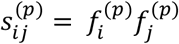. To distinguish N-glycosylation sites from other Ns in the sequence, we included a separate calculation for Ns in the NxS/T motifs. Focusing on aligned positions with conserved N (frequency of N >= 50%), we calculated the overall substitution frequency between amino acids *i* and *j* as 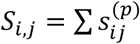, and substitution rate from N (or Nx/ST) to any amino acid *a* was calculated as *R*_*N,a*_=*S*_*N,a*_/Σ*S*_*N,i*_ and *R*_*N,a*_ =*S*_*NxST,a*_/Σ*S*_*NxST,i*_

## Supplemental Figures and Tables

**Supplemental Figure 1.**
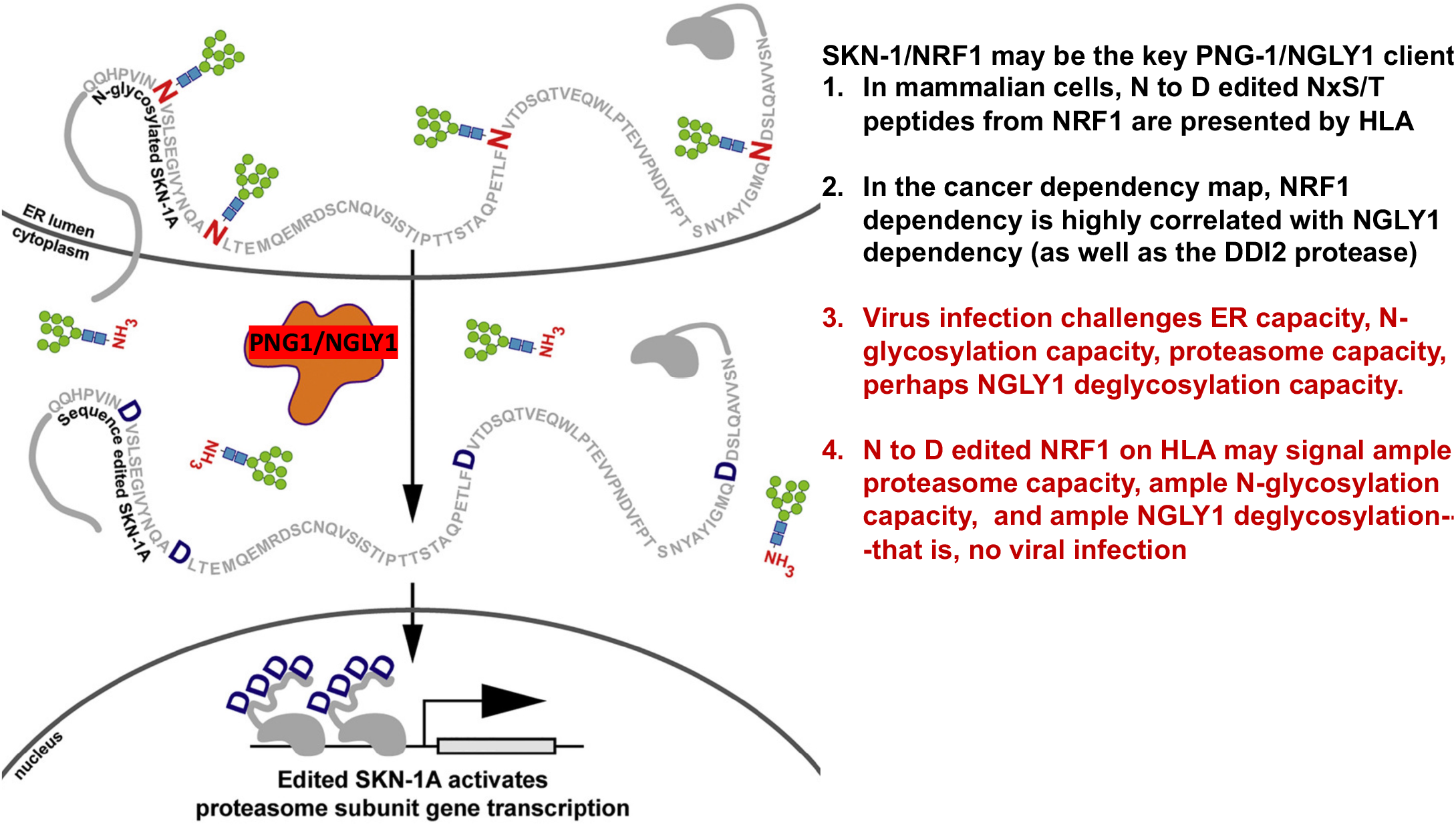
Summary of NGLY1/PNG-1 editing of SKN-1A/NRF1 as signal of normal N-glycosylation and deglycosylation function.

**Supplemental Figure 2.**
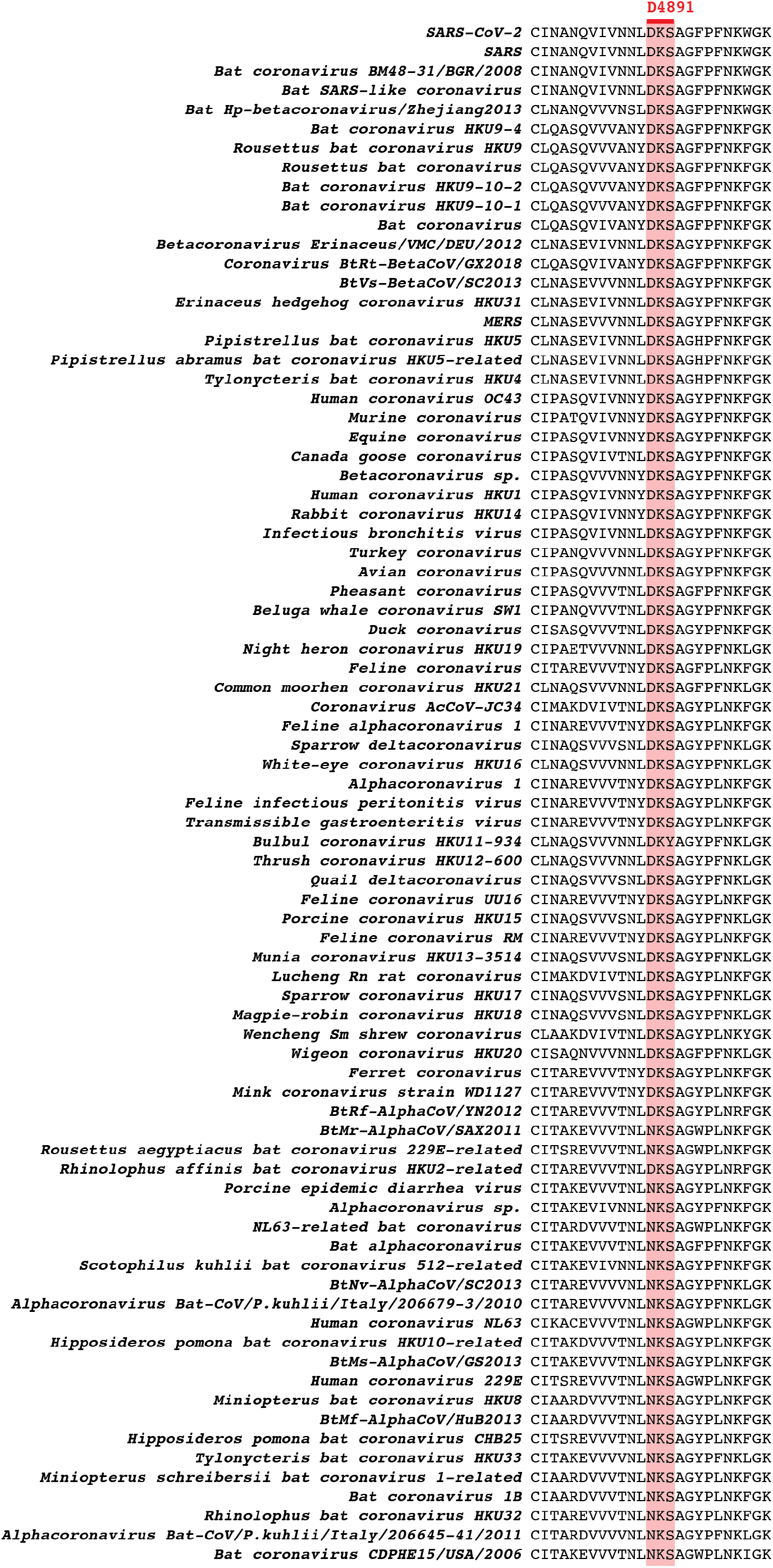
Multiple sequence alignment in the vicinity of N-glycosylation site in the RdRP epitope of human coronavirus NL63 that shows the N to D substitution in SARS-Cov2 and many other coronaviruses.

**Supplemental Figure 3.**
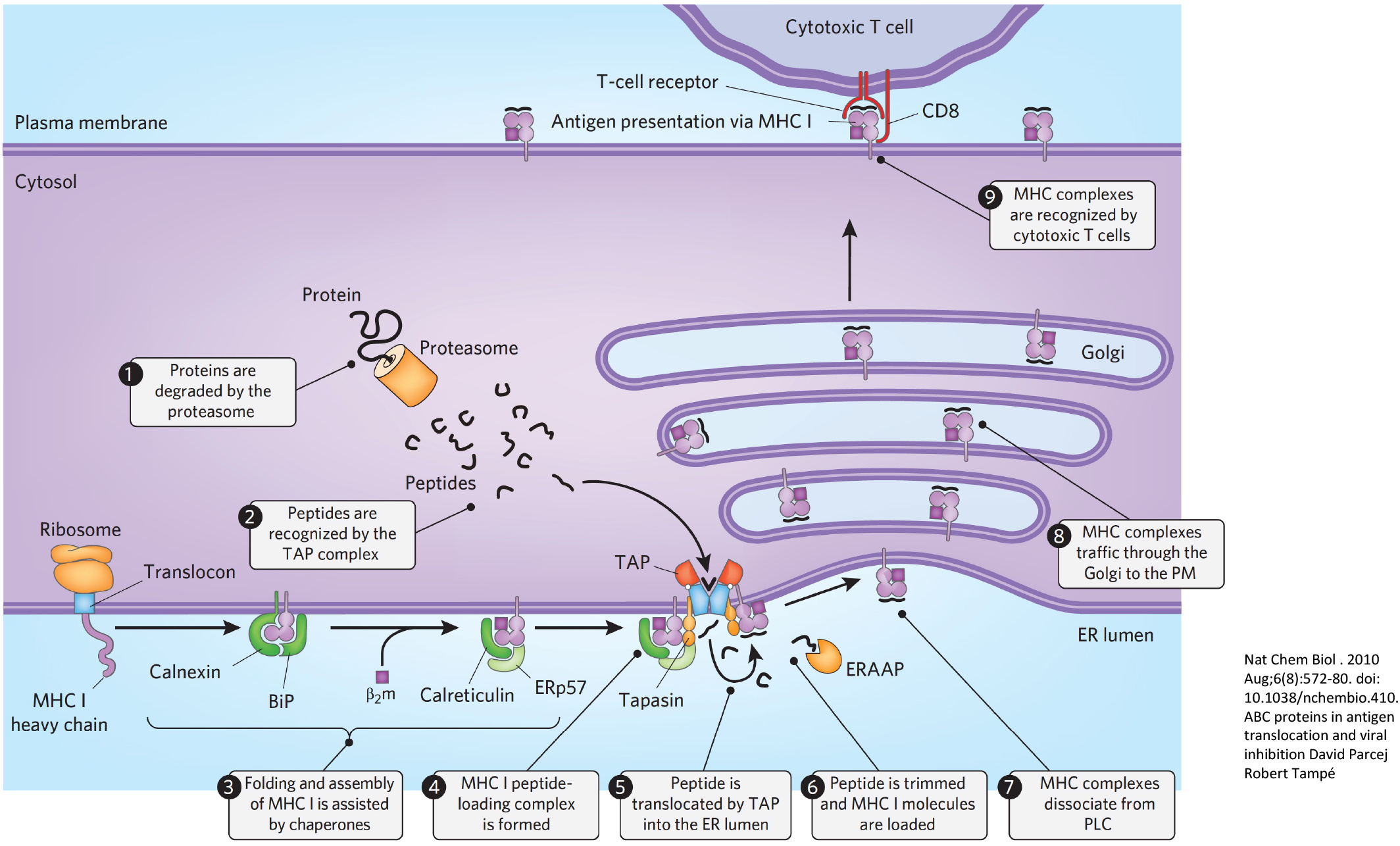
Steps in MHC loading of peptide fragments from the Proteasome. From Parcej and Tampe.

**Supplemental Table 1:**
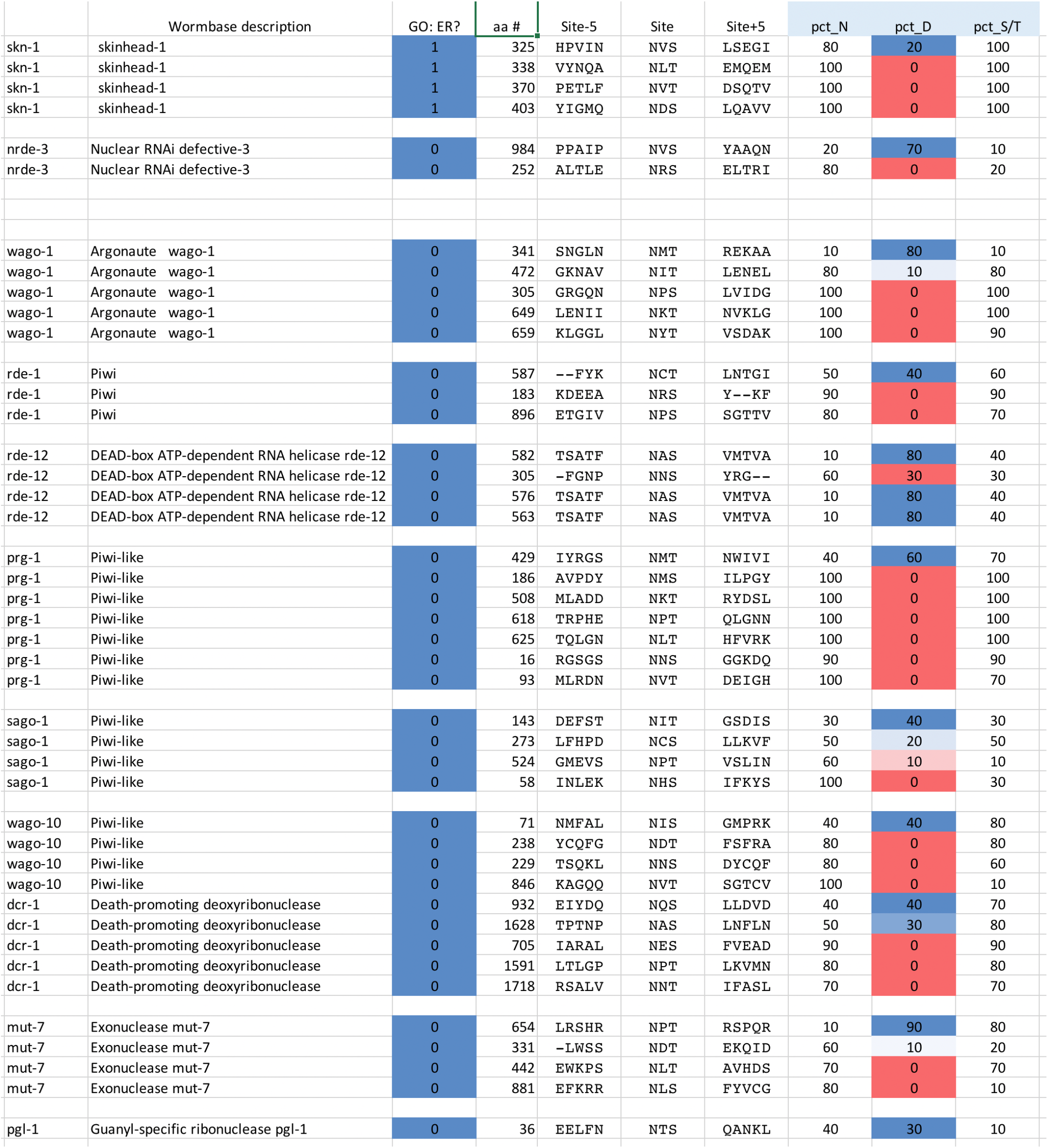
N to D variation between C. elegans and other Caenorhabditae in SKN-1A, Argonaute proteins and other antiviral RNAi factors. Some examples of *C. elegans* NxS/T sites that show evidence of substitutions to D in 10% to 90% of related Caenorhabditis species. Shown are the protein sequences at position -5 of the NxS/T site, the 3 amino acids of the NxS/T site, and the protein sequences at +5 from the NxS/T site. The percentage of N or percentage of D at position 1, as well as the conservation of S or T at position 3 are also shown, as are the 13 amino acid segments in each Caenorhabditis species.

**Supplemental Table 2 (an Excel spreadsheet as a separate file)**

Shown are the Virscan peptides from RNA viruses including many coronaviruses that show a frequency of N to D substitution in NxS/T sites of the query virus. For some segments of the viruses, the query protein can be aligned with hundreds of diverse non-identical viruses, whereas for some proteins, a number as small as 37 other viruses can be aligned (shown in column K). Sorted by percentage of D substitions. For those viruses, with 97 percent D for example, the query virus with an NxS/T site is the rare version, with the DxS/T much more common.

